# Inhibitory regulation of calcium transients in prefrontal dendritic spines is compromised by a nonsense *Shank3* mutation

**DOI:** 10.1101/2020.01.07.894832

**Authors:** Farhan Ali, Ling-Xiao Shao, Danielle M. Gerhard, Katherine Sweasy, Santosh Pothula, Christopher Pittenger, Ronald S. Duman, Alex C. Kwan

## Abstract

The *SHANK3* gene encodes a postsynaptic scaffold protein in excitatory synapses, and its disruption is implicated in neurodevelopmental disorders such as Phelan-McDermid syndrome, autism spectrum disorder, and schizophrenia. Most studies of SHANK3 in the neocortex and hippocampus have focused on disturbances in pyramidal neurons. However, GABAergic interneurons likewise receive excitatory inputs and presumably would also be a target of constitutive SHANK3 perturbations. In this study, we characterize the prefrontal cortical microcircuit in awake mice using subcellular-resolution two-photon microscopy. We focused on a nonsense R1117X mutation, which leads to truncated SHANK3 and has been linked previously to cortical dysfunction. We find that R1117X mutants have abnormally elevated calcium transients in apical dendritic spines. The synaptic calcium dysregulation is due to a loss of dendritic inhibition via decreased NMDAR currents and reduced firing of dendrite-targeting somatostatin-expressing (SST) GABAergic interneurons. Notably, upregulation of the NMDAR subunit GluN2B in SST interneurons corrects the excessive synaptic calcium signals and ameliorates learning deficits in R1117X mutants. These findings reveal dendrite-targeting interneurons, and more broadly the inhibitory control of dendritic spines, as a key microcircuit mechanism compromised by the SHANK3 dysfunction.

## Introduction

SHANK3 is a scaffold protein that is localized to the postsynaptic membrane, and supports the formation, modification, and maintenance of glutamatergic synapses [1, 2]. Disruptions of the *SHANK3* gene have been associated with multiple neurodevelopmental disorders [3]. Loss-of-function mutations in *SHANK3* cause the 22q13.3 deletion disease, also known as Phelan-McDermid syndrome [4]. Signs appear early in postnatal development, and commonly include cognitive delays, absent speech, low muscle tone, and seizures. Furthermore, the same individuals frequently have intellectual disability and autistic features. Studies have brought forth evidence linking *SHANK3* to these disorders by finding mutations and copy number variants in individuals with autism spectrum disorder [5, 6], which is confirmed by an extensive screen and meta-analysis [7]. Other clues suggest that certain *SHANK3* variants may contribute to the etiology of schizophrenia [8]. In particular, R1117X is a *de novo* nonsense mutation in *SHANK3* in three affected brothers identified during a screen of schizophrenia patients [9]. Interestingly, the brothers did not have dysmorphic features or repeated seizures that are typical with the aforementioned disorders, but instead were diagnosed with adolescent-onset schizophrenia or schizoaffective disorder. Therefore, a spectrum of behavioral phenotypes can arise from *SHANK3* mutations, presumably due to their varying impacts on the brain circuitry, which are not understood.

Studies of SHANK3 in animal models have mostly relied on gene deletion such as mice with deletion of one or several major isoforms of *Shank3* [10-13], and primates with deleterious mutations introduced by genome editing [14]. These studies have revealed dysfunction in distributed brain regions including the striatum, hippocampus, and neocortex. For the hippocampus in mouse models, pyramidal neurons are reported to have diminished synaptic function, measured as reduced post-tetanic potentiation in adults [12], and deficient glutamatergic transmission and reduced long-term potentiation in adolescents [13]. The altered hippocampal transmission characteristics have been linked to a decrease in N-methyl-D-aspartate receptor (NMDAR)-mediated synaptic responses [15]. For the neocortex, specifically the prefrontal cortical regions, pyramidal neurons have reduced dendritic spine density and fewer synaptic receptors in a non-human primate model [14]. Likewise, in mouse models with *Shank3* deletion, prefrontal neurons exhibit a reduction in postsynaptic responses mediated by glutamatergic receptors including AMPARs [16] and NMDARs [17, 18]. These synaptic defects may underlie the reduced connectivity within the local prefrontal cortical network as well as in long-range cortico-cortical pathways [19]. Collectively, the reduced glutamatergic signaling in pyramidal neurons agrees with the expectation from losing a protein crucial for maintaining the integrity of the excitatory postsynaptic machinery.

Although most work on the SHANK protein family have emphasized the principal neurons, several studies have reported alterations related to minority cell types, such as interneurons. For example, in certain *Shank3* mutants, prefrontal pyramidal neurons have decreased frequency of spontaneous inhibitory synaptic events [20], suggesting reduced inhibitory control. Characterizations of interneurons have focused on the analyses of synaptic marker staining, showing fewer GABAergic contacts onto pyramidal neurons in related mouse models [21, 22]. Moreover, there are hints that drugs targeting the GABA-A receptor can normalize SHANK-related abnormalities in neural activity patterns or behavioral deficits [23, 24]. However, GABAergic interneurons are heterogeneous, and there are multiple major subtypes in the neocortex and hippocampus. These GABAergic subpopulations differ greatly in their function and morphology, most notably in their axonal targeting to inhibit specific compartments of the pyramidal neurons [25]. In particular, the somatostatin-expressing (SST) interneurons specialize in innervating dendrites and dendritic spines. The extent to which *Shank3* mutations may impact these dendrite-targeting GABAergic neurons and the functional consequences on synaptic responses are unknown.

To study how SHANK3 dysfunction impacts the cortical microcircuit, we focus on a nonsense R1117X mutation. Notably, mice harboring the R1117X mutation in *Shank3* have previously been shown to have disrupted synaptic function in the medial prefrontal cortex [26]. In adult R1117X mutants, prefrontal cortical neurons had fewer and smaller miniature excitatory postsynaptic currents, which accompanied a reduction in dendritic spine density. The cortical defects are accompanied by a constellation of behavioral abnormalities including social dominance in the form of aggressive barbering of cage mates, as well as reduced locomotor activity and increased anxiety. Intriguingly, these synaptic and behavioral alterations are unlike those found in counterpart mouse mutants with an ASD-linked loss-of-function mutation. However, initial characterizations of the synaptic properties for R1117X mutants have been limited to *in vitro* single-cell recordings, and it is difficult to extrapolate to *in vivo* conditions because cortical neurons are densely interconnected and embedded in microcircuits. Therefore, we expect an *in vivo* study of the R1117X mutation will provide unique insights into how a *Shank3* mutation disrupts neural circuit function and translates to distinct behavioral characteristics.

To characterize the impact of the R1117X mutation on synaptic function and cortical microcircuit *in vivo*, here we use two-photon microscopy to measure calcium transients in dendritic spines and GABAergic neurons in the medial prefrontal cortex of adolescent R1117X mutant mice. Contrary to the expectation of decreased calcium influx from reduced glutamatergic signaling, we surprisingly find an abnormal elevation of synaptic calcium signals in the apical dendritic spines of pyramidal neurons. To determine the mechanism, we image from SST GABAergic interneurons and find reduced activity corresponding to a loss of dendritic inhibition. Corroborating evidence for the microcircuit-level dysfunction came from causal experiments, because increasing dendritic inhibition by manipulating NMDAR signaling in SST interneurons is sufficient to normalize synaptic calcium signals and restore prefrontal cortex-dependent learning in R1117X mutants. Together, these results uncover specific *in vivo* synaptic and microcircuit deficits that could help explain the cortical dysfunction associated with the nonsense *Shank3* mutation.

## Materials and methods

### Animals

All experimental procedures were approved by the Institutional Animal Care and Use Committee at Yale University. To study the R1117X mutation in *Shank3*, we used homozygous males of the Shank3*R1117X knock-in mouse line [26] (Stock No. 028779, Jackson Laboratory). To target SST interneurons, we created double transgenic SST-Cre/R1117X^+/+^ mice by first mating homozygous R1117X^+/+^ mice with homozygous SST-IRES-Cre mice (henceforth, SST-Cre) [27] (Stock No. 013044, Jackson Laboratory) to create mice that are heterozygous at both loci. We paired up these mice for another round of breeding, and genotyped to select for SST-Cre/R1117X offspring that were homozygous for R1117X mutation and positive for a least one copy of Cre (SST-Cre/R1117X^+/+^; 3/16 of all offspring based on Mendelian inheritance). To study a different *Shank3* mutation, we used *Shank3B*^+/-^ mice (Stock No. 017688, Jackson Laboratory) with a global knockout of PDZ domain (exons 13-16) of the *Shank3* gene [10]. Wild type littermates were used as controls for R1117X^+/+^ and *Shank3B*^+/-^ experiments, and SST-Cre littermates were used as controls for SST-Cre/R1117X^+/+^ experiments. For imaging experiments, we used adolescent males that are 5 – 6 weeks of age at the start of behavioral or imaging experiments. For trace fear conditioning, we used 8 – 12-week-old males, because adolescent R1117X^+/+^ males had low locomotor activity in an operant chamber, leading to spurious detection of freezing behavior. Genotyping was done by real-time PCR (Transnetyx, Inc.). Mice were group housed (3-5 mice per cage) on a 12:12-hr light/dark cycle with free access to food and water. Experiments were typically done during the light cycle. Sample sizes for this study were determined based on numbers typically employed in the field. There was no randomization when animal subjects came from different strains.

### Surgery

For all stereotaxic surgical procedures, the mouse was anesthetized with isoflurane in oxygen (3-4% during induction, 1-1.5% for the remainder of the surgery), and placed in a stereotaxic apparatus (David Kopf Instruments). The animal lay on top of a water-circulating pad (Gaymar Stryker) set at a constant temperature of 38 °C. Eyes were lubricated with ophthalmic ointment. The mouse was injected with carprofen (5 mg/kg, s.c.; 024751, Henry Schein Animal Health) and dexamethasone (3 mg/kg, i.m.; 002459, Henry Schein Animal Health). The skin above the skull was sterilized by ethanol and betadine swabs before an incision was made to expose the skull. Using a dental drill, a small craniotomy (∼0.5 mm diameter) was made centered at 1.5 mm anterior of bregma (anterior-posterior, AP) and 0.3 mm lateral of midline (medial-lateral, ML) to target the medial prefrontal cortex (Cg1 and M2 regions) in the right hemisphere. A glass pipette (3-000-203-G/X, Drummond Scientific) was pulled to a fine tip (P-97 Flaming/Brown Micropipette Puller, Sutter Instrument), front-filled with the relevant viruses (see below) and then lowered into the brain at 4 sites corresponding to vertices of a 0.1-mm wide square centered at the above stereotaxic coordinates. At depth of 0.3 to 0.5 mm below the dura, we ejected the virus using a microinjector (Nanoject II, Drummond Scientific). For all injections, we waited at least 5 minutes after finishing injection at one site before retracting the pipette to inject in the next site so as to reduce backflow of the virus. For two-color imaging, we used a similar approach to target the retrosplenial cortex (RSC), centering at AP = -1.4 mm, and ML = 0.3 mm, and injected the viruses sequentially. The brain was irrigated with artificial cerebrospinal fluid (ACSF (in mM): 5 KCl, 5 HEPES, 135 NaCl, 1 MgCl^2^, 1.8 CaCl^2^; pH 7.3). After the final injection, the craniotomy was covered with silicone elastomer (0318, Smooth-On, Inc.), the skin was sutured (Henry Schein), and the animal was returned to its home cage. After approximately 1-2 weeks, the animal would undergo a second surgery for cranial window implant. After the initial steps of anesthesia as outlined above, a larger incision was made to remove all of the skin above the skull, which was further cleaned to remove connective tissues. A 3-mm diameter craniotomy was made around the previously targeted location. A glass window was made by gluing two 3-mm-diameter, #1-thickness circular glass (640720, Warner Instruments), using UV-sensitive optical adhesive (NOA61, Norland Products). The glass window was placed on the exposed brain with slight downward pressure while high-viscosity adhesive (Loctite 454) was applied around the perimeter of the glass window. After the adhesive cured, a custom-made stainless steel headplate (eMachineShop.com) was then affixed to the skull with quick adhesive cement (C&B Metabond, Parkell). Immediately after surgery and on each day for the following 3 days, carprofen (5 mg/kg, s.c.) was given to the animal. All animals had at least 1 week to recover before starting experiments unless stated otherwise. For behavioral experiments, there was no second surgery for cranial window and head plate implant, injections were bilateral, and we waited at least 4 weeks after surgery before behavioral testing.

### Immunofluorescence staining

Mice were transcardially perfused with paraformaldehyde solution (4% (v/v) in phosphate-buffered saline (PBS)). The brains stayed in the fixative for at least 48 hours, and then they were sectioned at 100-μm thickness with a vibratome. For free-floating immunofluorescence staining of SST interneurons, sections were washed with PBS for 3 times (1 – 2 min each) in between all change of reagents. Sections containing Cg1 and M2 were first incubated in citrate buffer (diluted from 10x citrate buffer, pH 6.0; ab64214, Abcam) for 45 min at 95°C. After cooling at room temperature (RT), sections were incubated for 1 hr in blocking solution (PBS with 0.5% Triton X-100 and 5% goat serum) at RT followed by rat monoclonal anti-SST primary antibody (1:100; MAB354, Sigma-Aldrich) for at least 12 hr at 4°C. The specificity of the primary antibody was previously shown using preadsorption of synthetic SST peptide [28]. Sections were then incubated in secondary antibody for 3 hr at RT. We used secondary antibodies conjugated to different fluorophores depending on the fluorescent proteins in the brain tissue. For quantifying double-labeling with GCaMP6s, we used goat anti-rat IgG H&L Alexa Fluor 555 (1:500; ab150158, Abcam). For quantifying triple-labeling with mTagBFP2 and tdTomato, we used goat anti-rat IgG H&L Alexa Fluor 488 (1:500; ab150157, Abcam). The sections were then washed with filtered H_2_O before being mounted on slides, air-dried in the dark, and sealed with coverslips using DPX mounting medium (06522, MilliporeSigma) for long-term storage. The sections were then imaged with an upright fluorescence microscope (Zeiss Axio Imager M2 or Olympus BX61). Initially we would identify the central injection location by determining the two adjacent Cg1/M2 coronal sections with the largest extent of labeling. From these coronal sections, five random fields of view (20X; 512 × 512 μm) in the center of the injected region were then taken per animal. Cells labeled by the respective fluorophores were counted to quantify the specificity and efficiency of the viruses.

### Viruses

To visualize calcium transients in pyramidal neurons and their dendrites, we injected AAV1-CamKII-GCaMP6f-WPRE-SV40 (Penn Vector Core) diluted to a titer of 2 × 10^12^ genome copies (GC) per milliliter. A total of 73.6 nL of virus was injected for each experiment. To image calcium transients in SST interneurons, we injected AAV1-Syn-DIO-GCaMP6s-WPRE-SV40 (Penn Vector Core; 1 × 10^12^ GC/mL titer, 147.2 nL) in SST-Cre or SST-Cre/R1117X^+/+^ animals. For the two-color imaging experiment, two viruses were injected in the same animal: AAV1-Syn-NES-jRGECO1a-WPRE-SV40 (Addgene; 1 × 10^13^ GC/mL titer, 294.4 nL) in Cg1/M2, and AAV1-Syn-GCaMP6s-WPRE-SV40 (Penn Vector Core; 3 × 10^13^ GC/mL titer, 294.4 nL) in RSC. To upregulate NMDAR expression in a cell-type-specific manner, mouse GluN1 cDNA (OMu21895D, Genescript) and mouse GluN2B cDNA (EX-Mm24581-M02, Genecopoeia) were cloned to custom-made vectors. Other cassettes were either cloned from other existing vectors or synthesized *de novo* (IDT DNA Technologies). The vectors were then packaged into lentiviruses pseudotyped with VSV-G to produce: LV-Syn-DIO-GluN1-2A-tdTomato (Vigene Biosciences, 2 × 10^9^ transducing units (TU)/mL) and LV-Syn-DIO-mTagBFP2-2A-GluN2B (SignaGen Laboratories, 1 × 10^9^ TU/mL). For control fluorophores, we injected AAV1-Syn-DIO-tdTomato (Addgene) and AAV9-CAG-DIO-mTagBFP2 (Vector Biolabs). For lentiviral manipulations, a total of 993.6 nL of each lentivirus or AAV was injected in Cg1/M2 for behavioral and slice electrophysiology experiments. The same amounts were injected in Cg1/M2 for imaging experiments followed by either AAV1-CamKII-GCaMP6f-WPRE-SV40 (in R1117X^+/+^ and wild type littermates for pyramidal neuron imaging) or AAV1-Syn-DIO-GCaMP6s-WPRE-SV40 (in SST-Cre/R1117X^+/+^ and SST-Cre littermates for SST interneuron imaging).

### Cell culture validation

We performed experiments with cell cultures to validate the viral-mediated manipulation of NMDAR expression. Primary neuronal cultures were prepared as previously described [29]. Briefly, cortical neurons at E18 from pregnant female rats were dissected, dissociated and plated in wells (∼1 million cells/well) and cultured in standard media. Viruses were added at DIV5 stage. To validate GluN1 overexpression, we used the following viruses: AAV1-CMV-Cre-GFP (1 × 10^13^ GC/ml titer, 400 nL) and LV-Syn-DIO-GluN1-2A-tdTomato (2 × 10^9^ TU/ml titer, 2 μL). Using these viruses, we made the four well conditions: GluN1, no Cre; GluN1, Cre; no GluN1, Cre; no GluN1, no Cre (no viruses added). After at least 21 days of culture, we performed Western blot for GluN1 (#5704S, Cell Signaling, 1:500, rabbit monoclonal antibody) and beta-actin (#4967S, Cell Signaling, 1:1000, rabbit polyclonal antibody) (**Supplementary Fig. 4a**). To validate GluN2B overexpression, we used identical procedures except that we used LV-Syn-DIO-mTagBFP2-2A-GluN2B (1 × 10^9^ TU/ml titer, 2 μL) and GluN2B antibody (#4207S, Cell Signaling, 1:1000, rabbit polyclonal antibody) (**Supplementary Fig. 4b**). Bands were quantified using Bio-Rad Image Lab software and raw band intensity values were normalized to beta-actin levels.

### Two-photon calcium imaging

We used a laser-scanning two-photon microscope (Movable Objective Microscope, Sutter Instrument) controlled by the ScanImage software [30]. Excitation source was a tunable Ti:Sapphire femotosecond laser (Chameleon Ultra II, Coherent), focused onto the brain with a water immersion objective (XLUMPLFLN, 20X/0.95 N.A., Olympus). The time-averaged laser power used during experiments was typically less than 100 mW after the objective. We used filters with the following center excitation (ex) and emission (em) wavelengths (Semrock or Chroma): GCaMP6s/f, 920 nm (ex) and 525 nm (em); EGFP, 920 nm (ex) and 525 nm (em); tdTomato and dsRed, 1040 nm (ex) and 605 nm (em); mTagBFP2, 820 nm (ex) and 442 nm (em). For two-color imaging, we excited GCaMP6s and jRGECO1a at 1000 nm, and collected emission around 525 nm and 605 nm for the two fluorophores respectively. Emitted photons were collected by a GaAsP photomultiplier tube. Images acquired were 256 × 256 pixels at 0.68 or 0.54 μm per pixel resolution for dendritic spines and axonal boutons or 1.35 μm per pixel resolution for cell bodies. Images were acquired using bidirectional scanning with a set of galvanometer-based scanners at a frame rate of 3.62 Hz. In a subset of experiments, we imaged using a resonant scanner (RESSCAN-MOM, Sutter Instrument) at a frame rate of 29.96 Hz. We used a behavioral control software program (Presentation, NeuroBehavioral Systems) to send a TTL pulse via a data acquisition device (USB-201, Measurement Computing) to the two-photon microscope at 30-s intervals, as well as to trigger all the other equipment. The Presentation software would save a log file containing timestamps such that data from imaging and other equipment (e.g., movements) can be synchronized in later analyses. Before the collection of any actual imaging data, mice were habituated to head fixation under the microscope for 3 - 5 days of increasing durations. During head fixation, the mouse would sit in an acrylic tube, which limits gross body movements but permits postural adjustments. We would image a field of view for 10 – 15 minutes. Occasionally, in the same animal, we would image a different field of view on a subsequent day. Imaging was done at 0 – 150 μm below dura for apical dendrites in layer 1, and 150 – 500 μm below dura for cell bodies and dendrites in layer 2/3. Based on the depth of the virus injection to target mostly layer 2/3 pyramidal neurons, the dendritic spines in layer 2/3 should include predominantly spines from basal dendrites, although we cannot rule out other types of dendrites (e.g., ascending dendrites from layer 5). We would image the medial portion of Cg1/M2, within 0 – 500 μm of the midline as assessed by the dark band of the midsagittal sinus. For imaging experiments involving cell-type-specific manipulation, during live imaging, we specifically targeted fields of view with co-labeled cells that expressed mTagBFP2 and tdTomato for overexpression experiment.

### Analysis of calcium imaging data

Starting from raw fluorescence images stored in multiple multipage tiff files, we first concatenated the files and then corrected for lateral motion using either TurboReg implemented as a plugin in ImageJ or NoRMCorre [31] in MATLAB. A custom graphical user interface (GUI) in MATLAB was developed to display the mean or variance projection of the fluorescence images, and facilitate manual selection of region of interests (ROI). The rest of the analysis depends on experiment type, and is as follows:

#### Dendritic spines

For dendritic spines, we manually scrolled through the imaging frames to look for ROIs that are likely single dendritic spines and dendritic shaft – neurite segments with multiple protrusions that display correlated patterns of fluorescence transients. The dendritic shaft ROIs were within 20 μm of the corresponding dendritic spine ROIs. Values of pixels within a ROI were averaged to generate *F*_ROI_(*t*). For each ROI, we would estimate the contribution of neuropil to the fluorescence signal. Specifically, we would calculate a radius, *r*, treating the ROI area as the area of a circle, and then creating an annulus-shaped neuropil ROI with inner and outer radii of 2*r* and 3*r* respectively. We excluded pixels if they belonged to the ROIs of other spines and shaft. We calculated the time-averaged signal for each pixel, and then determined the median for all pixels within the neuropil ROI. We excluded pixels if their time-averaged signal was higher than the median, effectively avoiding pixels belonging to unselected dendritic structures. We then averaged the time-dependent signal of non-excluded pixels in the neuropil ROI to generate *F*_neuropil_(*t*). To subtract the neuropil signal, we calculated

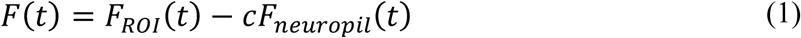

where *c* = 0.4 is the neuropil correction factor. The fractional change in fluorescence, Δ*F*/*F*(*t*), for each spine and shaft was calculated as follows:

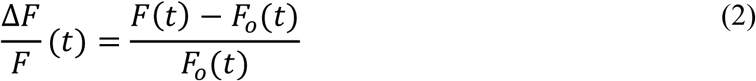

where *F*_0_ (*t*) is the 10^th^ percentile of *F*(*t*) within a 2-minute sliding window. Because we were mainly interested in calcium signals due to local synaptic inputs, we subtracted the calcium signals from non-local sources following a previously described procedure [32]. Briefly, for each spine, we computed the Δ*F*/*F*_synaptic_(*t*) by subtracting out the scaled version of fluorescence transients in the corresponding dendritic shaft, Δ*F*/*F*_shaft_(*t*), using the following equation:

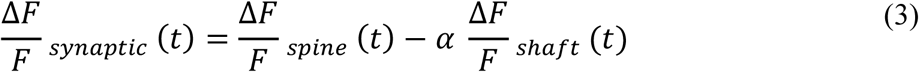

where α was determined by a linear regression forced through the origin of Δ*F*/*F*_spine_(*t*) versus Δ*F*/*F*_shaft_(*t*) (see an example in **Fig. 1c**).

**Fig. 1.**
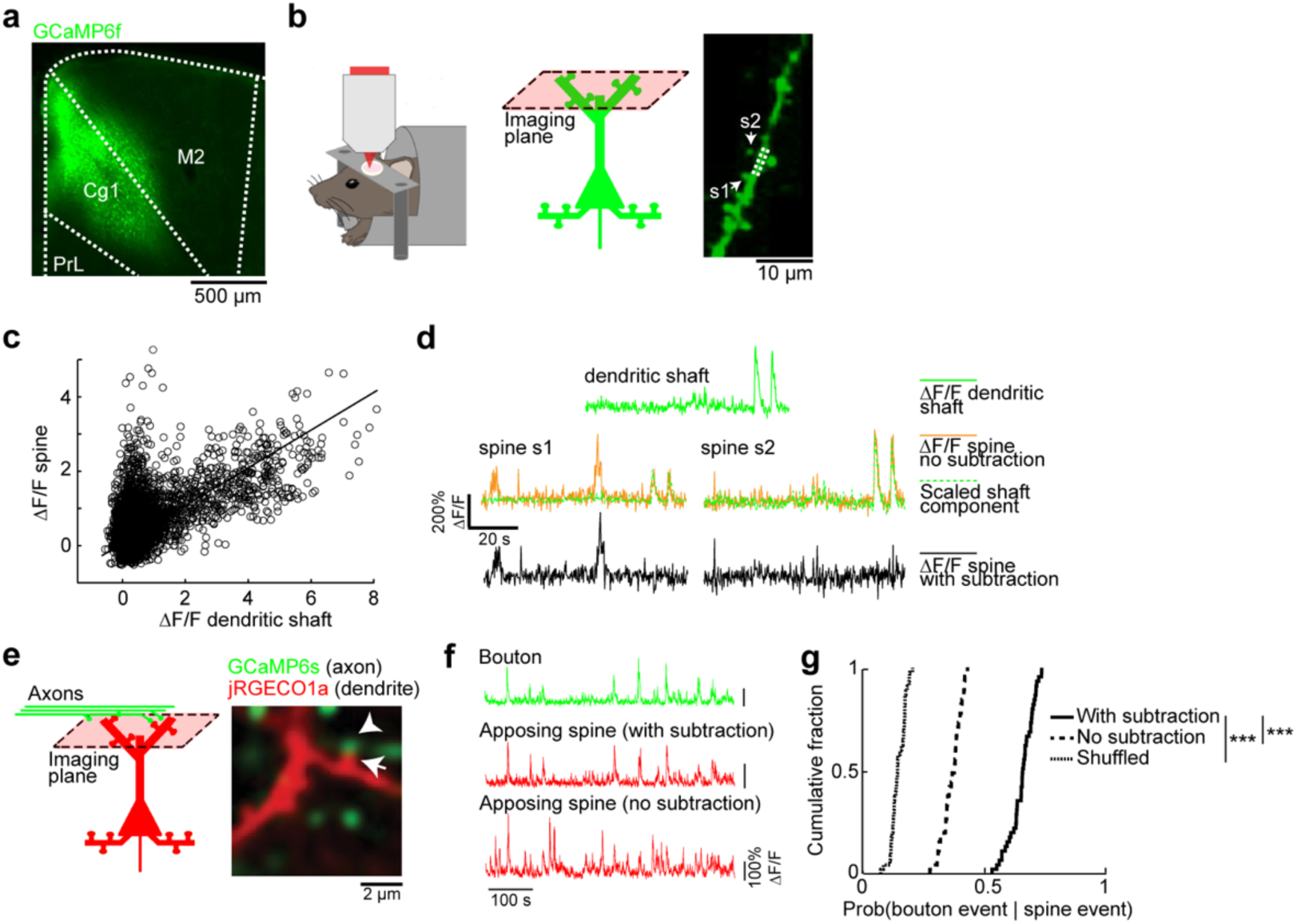
Spontaneous calcium transients in dendritic spines reflect subthreshold synaptic activation. (a) Coronal histological section, showing the extent of AAV-mediated expression of GCaMP6f in the mouse medial prefrontal cortex. Cg1, cingulate cortex. M2, secondary motor cortex. PrL, prelimbic cortex. (b) Left and middle, schematic of experimental setup and location. Right, an *in vivo* two-photon image of GCaMP6f-expressing apical dendritic spines in layer 1 of Cg1/M2 regions of medial prefrontal cortex. Two dendritic spines, s1 and s2, were identified based on their morphology as protrusions attached to the dendrite. Arrows, spines s1 and s2. Dashed lines, the dendritic shaft adjacent to the spines. (c) A scatter plot of the fluorescence transients (ΔF/F) measured from spine s1 against the ΔF/F measured from the adjacent dendritic shaft. Each open circle represents an image frame. Line, a least-squares regression line forced through the origin. (d) Top, example segment of ΔF/F signal recorded from the dendritic shaft (solid green line). Middle, ΔF/F signals recorded from the two dendritic spines, s1 and s2 (solid yellow lines), plotted along with the predicted non-local component of the spine calcium signal – the dendritic shaft calcium signal scaled based on the least-squares regression (broken green lines). Bottom, subtracting the scaled dendritic shaft ΔF/F signal from the dendritic spine ΔF/F signals to get an estimate of the synaptic calcium signals for spines s1 and s2 (solid black lines). (e) *In vivo* two-photon image of GCaMP6s-expressing long-range axons from retrosplenial cortex and jRGECO1a-expressing dendrites in the medial prefrontal cortex. Note a pair of apposing axonal bouton (arrowhead) and dendritic spine (arrow). (f) Fluorescence traces for the bouton and opposing spine (either with subtraction or no subtraction of dendritic shaft contribution) as denoted by the arrow and arrowhead respectively in (a). (g) Cumulative plot of the conditional probability of detecting a calcium event in an axonal bouton given detection of a calcium event in an apposing dendritic spine (data with subtraction: 0.65 ± 0.01, data without subtraction: 0.329 ± 0.004, shuffled: 0.144 ± 0.004, mean ± s.e.m.; *P* = 8 × 10^−43^, paired t-test). *n* = 45 bouton-spine pairs from 3 animals.

#### Axonal boutons

We selected ROIs that are likely single axonal boutons – varicosities on a relatively thin neurite branch with no protrusions. Importantly, the manual identification is corroborated by the known neuroanatomy. For SST interneurons, they send axons, but very few dendrites, if any, to layer 1 [33]. For retrosplenial cortical neurons, they send long-range axons to layer 1 of Cg1/M2. Values of pixels within an ROI were averaged to generate *F*_ROI_(*t*). For each ROI, we would estimate the contribution of neuropil to the fluorescence signal, by calculating and subtracting *F*_neuropil_(*t*) to obtain *F*_candidate_bouton_(*t*), using the procedures outlined in the last section. Some boutons may belong to the same neuron. To remove duplicates, we used a modified version of a previously published procedure [34]. Briefly, we first analyzed a subset of axonal imaging experiments to identify pairs of boutons on the same branch, based on distance (within 10 μm of each other) and visual observation of a connecting axonal segment. From this subset of bouton pairs (*n* = 46 bouton pairs, 2 animals), we used their *F*_candidate_bouton_(*t*) values to compute a correlation coefficient for each pair. The 5^th^ percentile value for all the correlation coefficients in this subset was 0.59, which was used as the cut-off threshold value. Then, for each axonal imaging experiment, we use an iterative procedure to identify unique boutons. In each iteration, we would compute a correlation matrix using *F*_candidate_bouton_(*t*) between all pairs of candidate boutons in a field of view. We would identify the pair of candidate boutons with the highest correlation coefficient, and proceed if the value exceeds the threshold of 0.59. We would randomly select one out of that pair of candidate boutons, and searched for other boutons whose correlation coefficient with the selected bouton exceeded the threshold of 0.59. The original identified pair and boutons from subsequent search then would belong to one cluster. Entries in the correlation matrix corresponding to this cluster would be excluded for subsequent iterations. The process iterated again to identify another cluster, and so on until no pair of candidate boutons in the correlation matrix had value above the threshold of 0.59. At the end of the iterative procedure, each of the remaining boutons was assigned to be its own cluster. For each cluster, we would randomly select one bouton as the representative. The fluorescence signals of these representative boutons, *F*_bouton_*(t)*, were then converted to fractional changes in fluorescence, Δ*F*/*F*_bouton_(*t*), and used for subsequent analyses.

#### Cell bodies

We selected ROIs that are likely cell bodies – fluorescent objects with a round boundary and a lateral extent of about 10 μm. Values of pixels within a ROI were averaged to generate *F*_ROI_(*t*). For each ROI, we would estimate the contribution of neuropil to the fluorescence signal, by calculating and subtracting *F*_neuropil_(*t*) to obtain *F*_soma_(*t*), and then calculating the fractional changes in fluorescence Δ*F*/*F*_soma_(*t*), using procedures outlined in the previous sections. For experiments in which we imaged a cell type with GluN2B manipulation, we analyzed only GCaMP6s-expressing cells that were co-labeled for both mTagBFP2 and tdTomato for GluN2B and GluN1 overexpression experiment.

#### Detection of calcium events based on template matching

After obtaining Δ*F*/*F*_synaptic_(*t*), Δ*F*/*F*_bouton_(*t*), or Δ*F*/*F*_soma_(*t*) for a ROI, we performed the following analysis to detect calcium events. We used a previously published “peeling” algorithm [35, 36], which employed an iterative, template-matching procedure to decompose Δ*F*/*F*(*t*) into a series of elementary calcium events. Briefly, we specified the template for an elementary calcium event to have an instantaneous onset with an amplitude of 30% Δ*F*/*F*(*t*), and a decay time constant of 1 s for a single-exponential function. The algorithm would search for a match to this template in Δ*F*/*F*(*t*). When a match occurred, a calcium event was noted, and the template subtracted from Δ*F*/*F*(*t*) (i.e., peeling), and the remaining Δ*F*/*F* trace was then searched for another match, in an iterative manner until no further matches were found. The output of the algorithm is the times of detected calcium events. The temporal resolution of the output is limited by the imaging frame rate. There can be multiple events associated with the same event time. In the original paper describing the peeling algorithm, extensive characterizations based on numerical simulations suggest excellent performance, even in worst-case scenarios with low signal-to-noise ratios and low sampling rate [36]. We nonetheless tested the robustness of our detection procedure. First, in a subset of data, we produced a reconstructed Δ*F*/*F* by convolving the template with a delta function at the detected calcium event times. We then computed correlation coefficient between the reconstructed Δ*F*/*F* and the recorded Δ*F*/*F* for all imaging frames with at least one detected calcium event (**Supplementary Fig. 1b**). This test demonstrated that the detected calcium events captured faithfully the measured fluorescence transients. Second, for the main results of this study as shown in **Fig. 2c**, we determined the extent to which differences between genotypes were sensitive to template parameters. We repeatedly analyzed the same data using detection procedures with a range of amplitude (30% – 150% in steps of 30%) and decay time constant (1 – 5 s in steps of 1 s) of the template (**Supplementary Fig. 1d**). This test demonstrated that, over a wide range of parameter values for detection procedure, the analysis would yield similar conclusions. For each imaging session, we calculated calcium event rate by dividing the number of calcium events by the duration of the imaging period (10 – 15 minutes). For **Fig. 2d, h, and i**, we characterized the calcium events in terms of amplitude and frequency. We identified calcium events that co-occurred in the same imaging frame, separating amplitude (mean number of calcium events per frame, for frames with at least one event) from frequency (number of frames with at least one event divided by the duration of the imaging session).

**Fig. 2.**
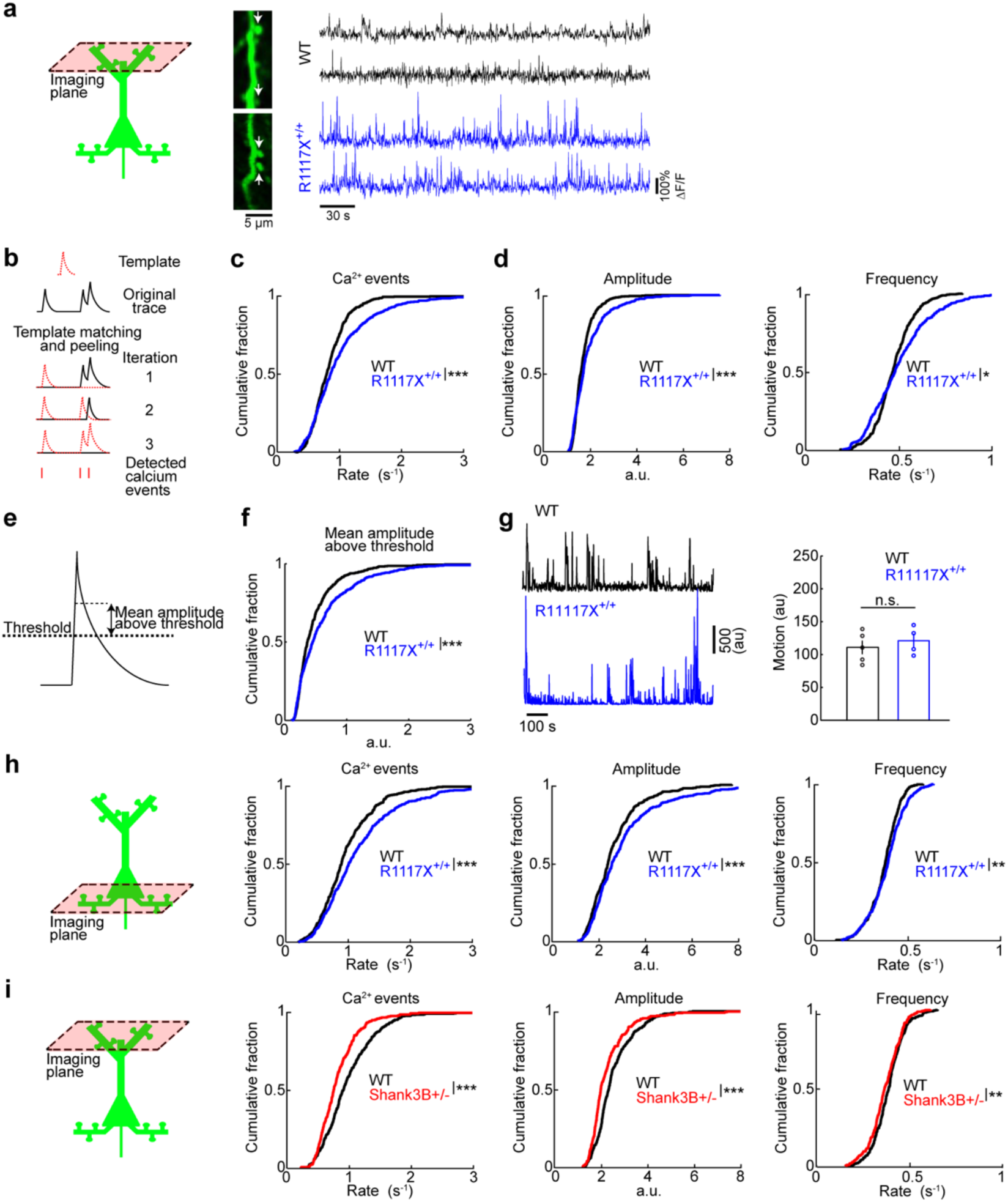
Aberrant calcium dynamics in apical dendritic spines in medial prefrontal cortex of R1117X mice. (a) Left, schematic of dendritic spine imaging in layer 1. Right, each row shows time-lapse fluorescence transients recorded from a dendritic spine. Two example spines came from wild type animals (black), and two other examples came from R1117X^+/+^ animals (blue). Locations of the spines in the *in vivo* two-photon images are indicated by white arrows. (b) Schematic illustrating the iterative template matching and peeling procedure to detect calcium events. (c) Cumulative plot of the rate of calcium events (R1117X^+/+^: 0.99 ± 0.02 Hz, wild type: 0.83 ± 0.01 Hz, mean ± s.e.m.; *P* = 6 × 10^−9^, two-sample t-test). For R1117X^+/+^, *n* = 553 dendritic spines from 4 animals. For wild type, *n* = 558 dendritic spines from 5 animals. (d) After a binning procedure (see Materials and methods), cumulative plots of amplitude (R1117X^+/+^: 2.57 ± 0.06, wild type: 2.22 ± 0.03, mean ± s.e.m.; *P* = 7 × 10^−8^, two-sample t-test), and frequency of binned calcium events (R1117X^+/+^: 0.390 ± 0.005 Hz, wild type: 0.376 ± 0.003 Hz, mean ± s.e.m.; *P* = 0.02, two-sample t-test). (e) Schematic illustrating a threshold-based procedure to detect calcium events. (f) Cumulative plot of mean amplitude above threshold (R1117X^+/+^: 0.62 ± 0.02; wild type: 0.48 ± 0.02; *P* = 7 × 10^−7^, two-sample t-test). (g) Left, example motion traces from a head-fixed animal during two-photon imaging session for wild type (black) and R1117X^+/+^ (blue). Right, time-averaged motion (mean ± s.e.m.) for wild type and R1117X^+/+^ (*P* = 0.6, Wilcoxon rank sum test). Each point is an imaging session. *n* = 5 animals for wild type, and 4 animals for R1117X^+/+^. (h) Schematic of dendritic spine imaging in layer 2/3. Cumulative plots of the rate of calcium events (R1117X^+/+^: 1.18 ± 0.03 Hz, wild type: 0.98 ± 0.02 Hz.; *P* = 3 × 10^−7^, two-sample t-test), as well as amplitude (R1117X^+/+^: 3.08 ± 0.09, wild type: 2.63 ± 0.05; *P* = 1 × 10^−5^, two-sample t-test) and frequency of binned calcium events (R1117X^+/+^: 0.388 ± 0.004 Hz, wild type: 0.371 ± 0.004 Hz.; *P* = 0.005, two-sample t-test). For R1117X^+/+^, *n* = 465 dendritic spines from 3 animals. For wild type, *n* = 419 dendritic spines from 4 animals. (i) Schematic of dendritic spine imaging in layer 1 of *Shank3B*^+/-^ animals. Cumulative plots of the rate of calcium events (*Shank3B*^+/-^: 0.85 ± 0.02 Hz, wild type: 1.015 ± 0.001 Hz.; *P* = 6 × 10^−6^, two-sample t-test), as well as amplitude (*Shank3B*^+/-^: 2.31 ± 0.06, wild type: 2.57 ± 0.05; *P* = 7 × 10^−4^, two-sample t-test) and frequency of binned calcium events (*Shank3B*^+/-^: 0.369 ± 0.005 Hz, wild type: 0.388 ± 0.004 Hz.; *P* = 0.003, two-sample t-test). * *P* < 0.05; ** *P* < 0.01; *** *P* < 0.001; n.s., not significant.

#### Detection of calcium events based on a threshold

In addition to the aforementioned template matching procedure, we used an alternative, threshold-based method to characterize calcium transients in dendritic spines. We started from Δ*F*/*F*_synaptic_(*t*). For each ROI and for each condition (i.e., pre- or post-injection), we calculated a threshold corresponding to 3 times the median absolute deviation of Δ*F*/*F*_synaptic_(*t*). We identified image frames in which Δ*F*/*F*_synaptic_(*t*) was above the threshold. From this subset of image frames, we determined the mean amplitude above threshold (Δ*F*/*F*_synaptic_(*t*) minus the threshold) (see **Fig. 2e** for an illustration).

#### Analysis of EGFP imaging

Imaging artifacts from animal motion could induce transients in the fluorescence signals. This issue is particularly relevant for small neuronal compartments such as dendritic spines and axonal boutons. To characterize motion-related signals in our setup, we injected AAV1-CMVIVS-DIO-EGFP (Vector Biolabs; 6 × 10^13^ GC/mL titer, 184 nL) into Cg1/M2 in adult male CamKIIa-Cre [37] (Stock No. 005359, Jackson Laboratory) and SST-IRES-Cre animals to image apical dendritic spines of excitatory neurons (*n* = 76 spines, 3 animals) and SST axonal boutons (*n* = 72 boutons, 3 animals) respectively. Imaging conditions were the same as our calcium imaging experiments. Analyses were the same, except there was no regression and subtraction of shaft signals for dendritic spines, and no clustering for axonal boutons. We compared the results of these control EGFP experiments to the GCaMP6f experiments to quantify the extent of motion artifacts in our data (**Supplementary Fig. 1a-c**).

#### Analysis of two-color imaging data

To confirm that Δ*F*/*F*_synaptic_(*t*) indeed captures the synaptic component of spine calcium transients, we performed two-color imaging in Cg1/M2, using the red calcium indicator jRGECO1a to label Cg1/M2 neurons and their dendritic spines, and the green calcium indicator GCaMP6s to label RSC neurons and their axonal boutons in Cg1/M2. Using our MATLAB GUI, we selected ROIs corresponding to dendritic spines and axonal boutons based on emission channel and morphology. The ROIs were drawn to be non-overlapping in order to mitigate any potential signal crosstalk. Apposing bouton-spine pairs were identified as those with a centroid-to-centroid distance of less than 0.8 μm. For each bouton-spine pair, using procedures described in earlier sections, we determined calcium events in the dendritic spine (including the subtraction steps) and axonal bouton. To obtain the conditional probability of observing a bouton event given a spine event (**Fig. 1g**), for each spine event, we asked whether there was at least one calcium event in the apposing bouton in the same frame, one frame before, or one frame after. As control, for each bouton-spine pair, we generated shuffled data by randomly permuting the series of frame-by-frame event counts, calculated the conditional probability as above and computed the mean over 100 replicates. Conditional probabilities for shuffled data were tabulated for all bouton-spine pairs to obtain the shuffled cumulative fraction of conditional probabilities in **Fig. 1g**.

### Recording animal movements during two-photon imaging

To determine the mouse’s movements during two-photon imaging (**Fig. 2g**), in a subset of experiments, we illuminated the animal with infrared light and obtained videos (640 × 480 pixels, 20Hz) of the animal’s face and body using an infrared camera (See3CAM_12CUNIR, e-con Systems). Although the animal was head-fixed, it could still move parts of its body including face, trunk, and limbs. To synchronize the infrared video with imaging, the infrared illuminator was triggered to turn off for 100 ms by the same TTL pulse used to trigger the two-photon microscope. In the analysis, the interval between dark frames therefore indicated a duration of 30 s. We computed motion *m* in 1-s bins:

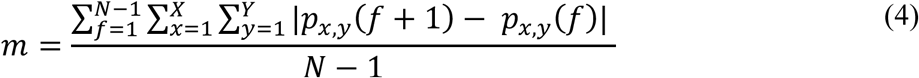

where *N* is the number of video frames in the 1-s bin, X and Y are the width and height of the frame, *p*_*x,y*_*(f+1)* and *p*_*x,y*_*(f)* are values of the pixel at coordinates *x* and *y* in frames *f+1* and *f* respectively.

### Cell-type-specific transcriptomics

To analyze cell-type-specific gene expression, we downloaded the dataset from Tasic et al. [38], available at the Allen Institute for Brain Science website. This was a dataset with single-cell transcriptomics from 1679 cells in the visual cortex of 8-week old mice which were subjected to clustering to identify cell classes and subclasses. Using the metadata of the downloaded dataset, we used the major class assignment “Excitatory” to group cells as excitatory neuron type. We used the subclass assignment of “Pvalb” and “Sst” to group cells as parvalbumin-expressing (PV) and SST interneuron types respectively. For all three neuron types, we combined data from all cortical layers. *Shank3* transcript level was then quantified for each of the three neuron types as reads per kilobase of transcript, per million mapped reads (RPKM).

### Slice electrophysiology

Three-week old animals underwent surgeries for viral injection in Cg1/M2 as described above. SST-Cre animals received AAVs, and SST-Cre/R1117X^+/+^ animals were randomly selected to receive lentiviruses or AAVs. The experimenter performing slice electrophysiology was blinded to the genotype and lentiviral manipulation. At about 6 weeks of age, brain slices from these animals were prepared for electrophysiology experiments. Under isoflurane anesthesia, mice were decapitated and 300-μm-thick coronal slices were cut in a vibratome (VT1000 S, Leica). The vibratome chamber was surrounded by ice and filled with a slicing solution containing (in mM): 110 choline, 25 NaHCO_3_, 1.25 NaH_2_PO_4_, 2.5 KCl, 11.6 sodium ascorbate, 7 MgCl_2_, 3.1 sodium pyruvate, 0.5 CaCl_2_, and 20 glucose. Three to four slices containing Cg1/M2 were then incubated in artificial cerebral spinal fluid (ACSF) containing (in mM): 127 NaCl, 25 NaHCO_3_, 1.25 NaH_2_PO_4_, 2.5 KCl, 1 MgCl_2_, 2 CaCl_2_ and 20 glucose. After incubation at 34°C for 30 min, the slices were maintained at room temperature and kept for at least 30 min before use. The slicing solution and ACSF were prepared every day with deionized water (18.2 MΩ-cm), filtered, and bubbled with 95% O_2_ and 5% CO_2_ for at least 15 minutes prior to use and throughout the slice preparation and recording experiment. At the beginning of each experiment, we placed the slice in an open diamond bath chamber that was perfused with ACSF at ∼3 ml/min and maintained at 34°C using an inline heater. For electrical stimulation, a tungsten bipolar electrode (2-3μm-diameter tip) with resistance of 0.5 MΩ (TST33A05KT, World Precision Instruments) was inserted into layer 2/3 of Cg1/M2. Use of a high-resistance, small-tip electrode reduced the occurrence of polyphasic current responses. For whole-cell patch clamp recordings, patch pipettes were pulled from borosilicate glass (BF-150-86-10, Sutter) to a resistance of 3 – 5 MΩ with a puller (P97, Sutter) and loaded with filtered internal solution containing (in mM): 125 CsMeSO_4_, 20 CsCl, 10 NaCl, 2 MgATP, 10 HEPES, 0.2 EGTA, 0.3 Na_3_GTP, 2.5 QX314 (pH 7.3, adjusted with CsOH). Under differential interference contrast (DIC) and fluorescence microscopy (BX51W, Olympus), we targeted two types of neurons in Cg1/M2. SST neurons were identified based on mTagBFP2 and tdTomato expression in SST-Cre or SST-Cre/R1117X^+/+^ mice. Putative pyramidal neurons were identified based on morphological criteria – large cell body and prominent apical dendrite – in wild type or R1117X^+/+^ mice. We recorded from cells that were approximately 200 μm away from the stimulating electrode. For all experiments, we added NBQX (20 μM) and picrotoxin (50 μM) to the ACSF to isolate NMDAR-mediated excitatory postsynaptic currents (NMDAR EPSC). In a subset of experiments, we bath-applied NMDAR antagonist, AP5 (50 μM), to confirm that the NMDAR EPSC was mediated by NMDARs. All recordings were made with a MultiClamp 700B amplifier (Molecular Devices) and digitized at 20 kHz (Digidata 1550, Molecular Devices). To determine evoked NMDAR response as a function of stimulation intensity, we set voltage clamp at +60 mV and applied stimulation intensities from 20 to 200 μA (single pulse, 0.5 ms duration) in steps of 20 μA using a stimulus isolator (Isoflex, A.M.P.I). Each stimulation intensity was given twice with an inter-stimulation interval of 20 s. To estimate the contribution of GluN2B, for each cell, we initially tuned the stimulation intensity to induce either 50 – 100 pA (SST neurons) or 200 – 250 pA (pyramidal neurons) of NMDAR EPSC at +60 mV. We recorded NMDAR EPSC before and after bath application of GluN2B-specific antagonist, ifenprodil (3 μM). Evoked responses from 8 – 20 repeats of the stimulation were obtained before and then again after ifenprodil. Because ifenprodil took time to wash out, we discarded the brain slice after each ifenprodil experiment, and would start again with a fresh brain slice. To characterize the activation curve, for each SST neuron, we initially tuned the stimulation intensity to induce 50 – 100 pA of NMDAR EPSC at +60 mV. We then clamped from -80 to +60 mV in steps of 20 mV. At each voltage step, we obtained the evoked response from 2 repeats of the stimulation. For analysis, we used Clampfit and MATLAB. Current traces were first processed by measuring the peak of NMDAR EPSC and subsequently averaged across repeats. At this point, the results were unblinded and the data collated according to genotype and manipulation.

### Behavior

#### Fear conditioning

Trace fear learning was assessed using a conditioning box equipped with an infrared video camera (320 × 240 pixels, 30 Hz; Med Associates, Inc.). To facilitate more flexible protocols involving multiple stimuli and repetitions, we modified the box to be controlled by a behavioral control software (Presentation, NeuroBehavioral Systems). Each of the conditioned stimuli (CS) was a 20-second long series of auditory pips (500 ms on, 500 ms off). The pip was either 2.5 kHz (85 dB, calibrated by A-weighting at 15 cm from speaker) or 11 kHz (75 dB). For delay conditioning experiments, one CS (CS+) co-terminated with the unconditioned stimulus of footshock (US; 0.65mA, 0.5 s), and the other CS (CS-) was not associated with any event. For trace conditioning experiments, the end of CS+ and US was separated by 15 s. Conditioning commenced after 6 minutes of habituation, after which 5 CS- and 5 CS+ were presented in a random order with a random inter-trial period of 100 – 110 s. The auditory characteristics of CS+ and CS- were counter-balanced across subjects. In the recall phase, which occurred about 24 hr later, animals were tested in a different context. Context was altered using different odors, wall inserts, light, and ambient noise. Testing commenced after 6 minutes of habituation, after which 4 CS- were presented followed by 4 CS+ (with no US) with a random inter-trial period of 100 – 110 s. Motion was assessed from the infrared video data using automated procedures in the Video Freeze software (Med Associates, Inc.). Briefly, according to the publication related to the software [39], motion was estimated by calculating the sum of pixel-by-pixel value changes across successive video frames. Time periods when the motion values fell below the software-preset threshold for at least 1 s were considered freezing periods. For 35 s after the onset of each CS+ or CS- (i.e., including the 20-second long CS presentation and 15-second long trace period), we determined the fraction of time in which freezing was detected. We reported the differential value, i.e., the fraction of time spent freezing in response to CS+, subtracted by the fraction of time spent freezing in response to CS-, as measured in the recall phase.

#### Prepulse inhibition

We used a commercial system (SR-Lab, San Diego Instruments) for prepulse inhibition as a measure of sensorimotor gating. An animal was placed in a cylinder on a platform with an accelerometer for recording motion in a soundproof chamber. For habituation, the animal was given 5 minutes of exposure to background noise (66 dB, calibrated by A-weighting at 25 cm from speaker) and startle stimuli (120 dB noise, 50 ms, 6 repeats). For testing, we presented 5 types of trials: startle, prepulse trials during which the onset of a prepulse stimulus (3, 6, or 9 dB noise above background, 50 ms) preceded a startle stimulus by 100 ms, and catch trials with no stimulus. A block consisted of 5 trials, 1 of each type, in a random order with a random inter-trial period of 8 – 23 s. For each animal, 10 blocks were presented (i.e., 50 trials in total). The prepulse inhibition (PPI) is calculated as follows:

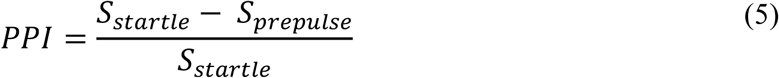

where *S*_startle_ and *S*_prepulse_ were the rectified motion signals recorded by the accelerometer during the 50-ms long startle stimulus averaged across trial repeats, for the startle and prepulse trials respectively.

#### Social dominance

To assess social dominance, we implemented the tube test assay [40]. A clear acrylic tube with inner diameter of 25 mm and length of 12 cm was used. The tube was wide enough to fit an adolescent mouse, but narrow enough to have no room for turning its body. A shortened version of the tube (length of 6 cm) was placed in the animal’s cage for up to 4 hours over two days to let the animals habituate to the tube in their home cage. Each mouse was then placed in a separate box to be further habituated over two further days to walking through the tube for at least 5 times on each side. Initially, mice were gently prodded to encourage a walk-through. On testing day, each mouse was paired with a stranger mouse matched for age and body weight. The stranger mouse was taken from a batch of ten SST-Cre mice that were distinct from the SST-Cre animals being tested. Each stranger mouse was used in no more than 4 matches. A match started when the two mice were grabbed by the tail, placed simultaneously on separate ends of the tube and gently pushed to the middle to face each other. The mouse that retreated by walking backward and exiting with four limbs outside the tube was considered the loser and the other mouse the winner. We repeated the match consisting of the same pair by counterbalancing for side of mouse entry. If a mouse in a pair won one match each, a third match was performed to break the tie. A time limit of 5 minutes was used. If no retreat was observed, the match was repeated. All repeats for the same pair were done at least 60 min apart on the same day.

#### Open field exploration

Animals were placed in a box (approx. 30 (width) x 25 (depth) x 20 (height) cm) with white ceiling lights. A camera (320 × 240 pixels, 30 Hz) was used to record the animal’s movements for 30 minutes. The video was then analyzed in MATLAB for time spent in the center of the box. We downsampled the video to 1 Hz and computed difference in pixel values for each pixel in neighboring frames. Because the biggest changes in pixel values corresponded to the animal instead of background noise, the algorithm was able to track the animal’s movement automatically on a frame-by-frame basis. We then computed the percentage of time the animal spent in the center over the whole 30 minutes. The center was defined as anything farther than 100 pixels from the corners, corresponding to a physical distance of approximately 9 cm.

#### Locomotion

To assess locomotor activity, we placed animals in the same box as for open field exploration for 30 minutes. Motion was assessed from the video data quantified using the same algorithm as the open field exploration. However, instead of finding the animal’s location, we simply summed the total difference in pixel values for each pixel in neighboring frames as measure of the overall movement of the animal. We averaged the movement trace in 1-min bins to plot the results for **Fig. 6f**. We then took the averaged across all 1-min bins for each animal as a measure of movement for comparison across genotypes.

### Statistics

Statistical tests as indicated in figure captions were performed in MATLAB. For the imaging experiments, all planned two-group comparisons were done with t-tests (two-tailed, α = 0.05). The sampling distribution of the mean was assumed to be normal, but this was not formally tested. For behavior and electrophysiology experiments, we used non-parametric tests (two-tailed, α = 0.05) except for experiments with multiple factors in which case we used ANOVA tests, with groupings as indicated in the figure caption, followed by post-hoc tests using Tukey-Kramer method if we had a statistically significant main or interaction effect. In the figures, *P*-values were represented as n.s., not significant; *P* > 0.05; \**P* < 0.05; \*\**P* < 0.01; \*\*\**P* < 0.001. See Figure captions for exact *P*-values.

## Results

### Characterizing calcium transients associated with subthreshold activation of dendritic spines

We injected AAV1-CamKII-GCaMP6f-WPRE-SV40 to express the calcium-sensitive fluorescent protein GCaMP6f in pyramidal neurons in Cg1/M2 (**Fig. 1a**), which constitutes the dorsal portion of the medial prefrontal cortex in rodents [41, 42]. Using a two-photon microscope, we imaged spontaneous calcium transients in the apical dendrites in layer 1 (<150 μm from the dura) in awake, head-fixed C57BL/6J wild type mice (**Fig. 1b)**. Previous studies have established that, upon a subthreshold synaptic input, calcium enters the spine compartment locally with minimal flow to the dendritic shaft [43, 44]. By contrast, more widespread regenerative events are expected to elevate calcium in both the shaft and spines. To focus on the synaptically driven component, we used a regression and subtraction analysis previously used by several studies [32, 45-47], which removes fluorescent transients that co-occur in the shaft and spine (**Fig. 1c, d**). The subtraction-isolated, local transients may then be quantified in terms of calcium events by using a ‘peeling’ algorithm to count transients that match an elementary template [36] (**Supplementary Fig. 1;** see also **Fig. 2b** for illustration). For the rest of the paper, the data presented are the subtraction-isolated local calcium transients determined using this method.

The analysis procedure is expected to isolate subthreshold synaptic calcium transients at a dendritic spine, but how well does it work? To determine empirically the effectiveness of our analysis procedure, we performed two-color calcium imaging in Cg1/M2, monitoring dendritic spines with the red-shifted indicator jRGECO1a [48], and axonal boutons from the retrosplenial cortex (RSC) with GCaMP6s (**Fig. 1e**). The chance of observing a jRGECO1a-expressing spine in Cg1/M2 with an apposing GCaMP6s-expressing bouton from RSC was low; typically, there were 1 – 2 spine-bouton pairs in each field of view that contains 30 – 50 spines. We analyzed specifically the few labeled spine-bouton pairs. Bouton fluorescent signals were correlated with the subtraction-isolated local spine fluorescence transients, and less so with the raw spine signals (**Fig. 1f**). Across the bouton-spine pairs examined, 65 ± 1% of calcium events in a spine were coincident with calcium events in the apposing bouton (mean ± s.e.m), compared to 32.9 ± 0.4% without subtraction and 14.4 ± 0.4% in shuffled controls (**Fig. 1g**). These results suggest that, for the spontaneously occurring signals in awake animals, the majority of the local calcium transients can be attributed to synaptic activation.

### Aberrant calcium dynamics in prefrontal dendritic spines of R1117X mice

For the R1117X mutants, we have focused the measurements on adolescent (5 – 6 weeks old), homozygous animals (R1117X^+/+^). Adolescence represents the time of onset for the symptoms associated with the mutation in the initial human study [9]. The use of homozygous animals was to accentuate the cortical defects, because the original report of the R1117X mice have characterized both heterozygous and homozygous mutants, and found that morphological, electrophysiological, and behavioral deficits occurred in a graded manner for the genotypes [26]. Imaging spontaneous calcium transients *in vivo* in prefrontal apical dendrites in layer 1, we found that the rate of synaptic calcium events was elevated in R1117X^+/+^ mutants relative to wild type littermates (**Fig. 2a-c**) (R1117X^+/+^: 0.99 ± 0.02 Hz, wild type: 0.83 ± 0.01 Hz, mean ± s.e.m.; *P* = 6 × 10^−9^, two-sample t-test;). To characterize the temporal dynamics of the aberrant transients, we identified calcium events that co-occurred in each image frame, thereby separating amplitude (number of events in each frame) from frequency (number of frames with at least one event). Both the amplitude and frequency parameters were elevated for the apical dendritic spines in layer 1 of R1117X^+/+^ mice (**Fig. 2d**). The result was insensitive to the choice of template parameters for event detection (**Supplementary Fig. 1d**). An alternative threshold-based event detection method yielded the same conclusion (**Fig. 2e, f**). The result held when the imaging was done at a higher frame rate (**Supplementary Fig. 2**). The result was also not due to motion artifacts because when imaging EGFP in dendrites, very low levels of false positive events were detected by our analysis procedures (**Supplementary Fig. 1a-c**). Moreover, the mutant and control animals did not have detectable differences in movements during imaging (**Fig. 2g**). Overall, these results revealed an abnormal elevation of synaptic calcium transients in the medial prefrontal cortex of R1117X^+/+^ mice.

To gain insight into the extent of the perturbation in pyramidal neurons, we imaged other cellular compartments including deeper-lying dendrites and cell bodies. In a different set of R1117X^+/+^ mice, we imaged dendritic spines in layer 2/3, presumably coming from basal dendrites, and found similar elevations in synaptic calcium transients (**Fig. 2h**). Cell bodies of layer 2/3 pyramidal neurons also exhibited higher levels of spontaneous calcium transients (**Supplementary Fig. 3**). To test whether the alteration in synaptic calcium signals was specific to the R1117X mutation in *Shank3*, we repeated the experiment in mice with a global knockout of the PDZ domain (exons 13-16) of the *Shank3* gene [10]. In contrast to the R1117X^+/+^ mice, the prefrontal dendritic spines in *Shank3B*^+/-^ animals had reduced rates of spontaneous calcium transients (**Fig. 2i**). This result suggests that different alterations to *Shank3* can lead to distinct functional consequences for the prefrontal dendritic spines.

### Diminished SST GABAergic interneuron activity *in vivo* in R1117X mice

The R1117X mutation leads to a truncated SHANK3 protein when tested in zebrafish and mice [9, 26]. As SHANK3 is a postsynaptic scaffold, any reduction of its function in pyramidal neurons should impair excitatory synaptic transmission, and thus lower synaptically evoked calcium influx. Instead, we observed the opposite effect in the prefrontal dendritic spines of R1117X mutants. Because the R1117X mutation is constitutive in the R1117X^+/+^ mouse, the mutation could affect other cortical cell types. Our analysis of data pulled from a single-cell transcriptomics database [38] indicated high levels of *Shank3* transcripts not only in cortical excitatory neurons, but also in cortical SST interneurons (**Fig. 3a**). SST interneurons are GABAergic cells that selectively inhibit dendritic spines and shafts of pyramidal neurons [49], and have been shown to powerfully modulate calcium influx in dendritic spines [50, 51]. Therefore, it is possible that the calcium dysregulation in dendritic spines stems from a microcircuit-level mechanism involving SST interneurons. To test the possibility, we determined cell-type specific activity by creating double transgenic SST-Cre/R1117X^+/+^ mice using SST-IRES-Cre animals [27] and expressing GCaMP6s in SST interneurons in the medial prefrontal cortex (Cg1/M2) (**Fig. 3b**). We verified the specificity and efficiency of GCaMP6s expression in SST interneurons (**Fig. 3c**). Compared to SST-Cre littermates, calcium event rates in SST cell bodies of SST-Cre/R1117X^+/+^ animals were markedly reduced (SST-Cre/R1117X^+/+^: 0.97 ± 0.08 Hz, littermates: 1.7 ± 0.1 Hz, mean ± s.e.m.; *P* = 1 × 10^−6^, two-sample t-test; **Fig. 3d, e**). Because SST cells are heterogeneous [52], to more firmly show that dendritic inhibition is reduced, we imaged those SST axonal boutons that would be innervating apical dendritic compartments in the superficial layers, and likewise found fewer calcium events in SST-Cre/R1117X^+/+^ mutants (**Fig. 3f**). These results indicate a loss of SST interneuron activity, which is expected to weaken dendritic inhibition in animals with the R1117X mutation.

**Fig. 3.**
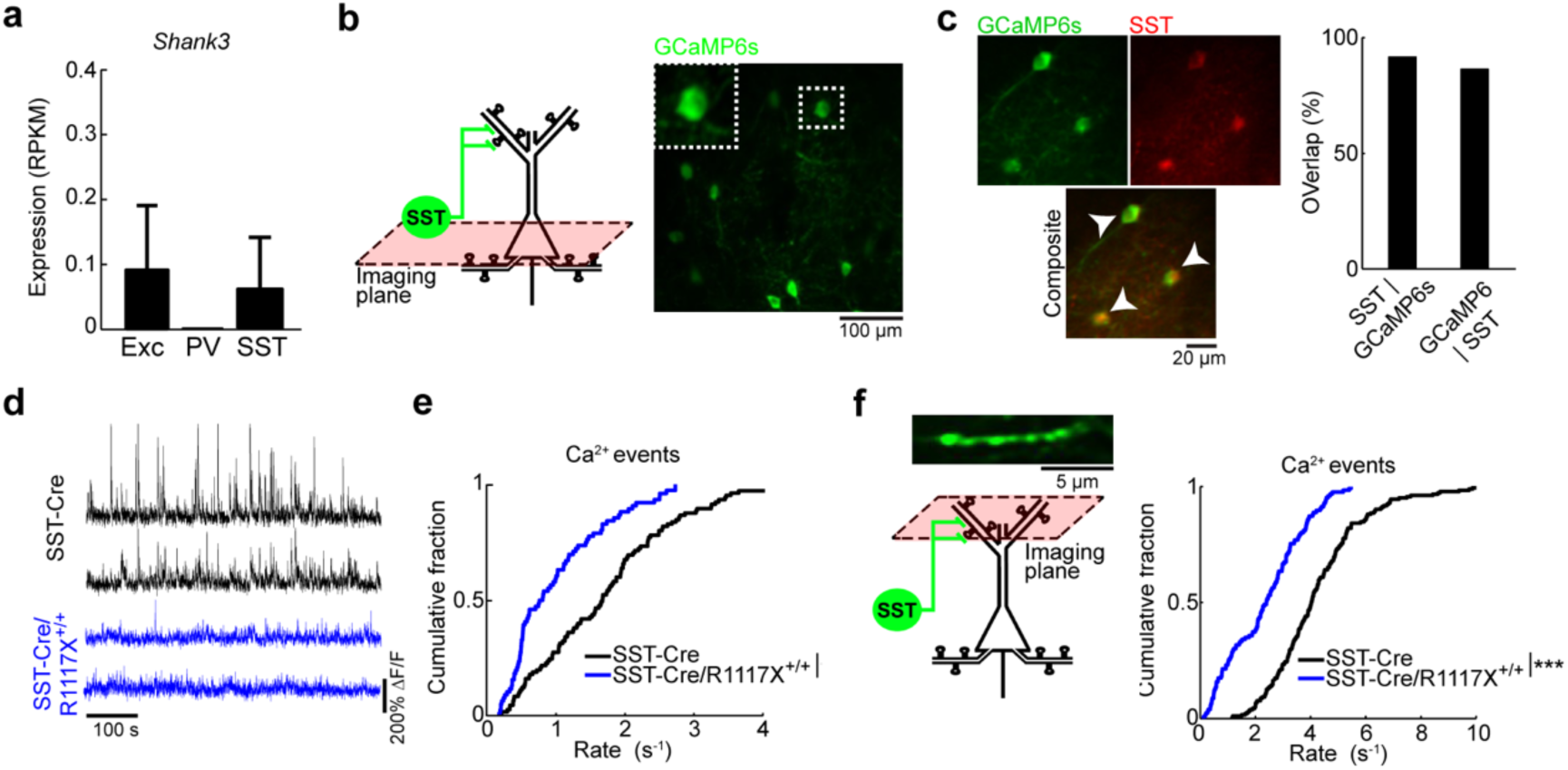
Reduced SST interneuron activity *in vivo* in R1117X mice. (a) Expression of *Shank3* transcripts in excitatory, PV, and SST neurons based on an analysis of the Allen Institute for Brain Science single-cell transcriptomics dataset. (b) Schematic of imaging location in layer 2/3, and an *in vivo* two-photon image of GCaMP6s-expressing SST interneurons in the medial prefrontal cortex (Cg1/M2) of an SST-Cre/R1117X^+/+^ animal. Inset, magnified view of the cell body of one SST interneuron. (c) Left, example images from post hoc immunofluorescence staining using an antibody against SST. Arrowheads, double-labeled cells. Right, quantification of specificity (SST | GCaMP6s, 91%) and efficiency of AAV (GCaMP6s | SST, 86%). *n* = 244 cells, 3 mice. (d) Each row shows time-lapse fluorescence transients from the cell body of one SST interneuron. Two example SST interneurons were plotted for SST-Cre littermates (black), and two other examples were plotted for SST-Cre/R1117X^+/+^ (blue). (e) Cumulative plot of the rate of calcium events for SST interneurons (SST-Cre/R1117X^+/+^: 0.97 ± 0.08 Hz, SST-Cre: 1.7 ± 0.1 Hz, mean ± s.e.m.; *P* = 1 × 10^−6^, two-sample t-test). For SST-Cre/R1117X^+/+^, *n* = 76 cells from 6 animals. For SST-Cre, *n* = 105 cells from 5 animals. (f) Same as (b, e), but for SST axons in superficial layers (SST-Cre/R1117X^+/+^: 2.4 ± 0.1 Hz, SST-Cre: 4.3 ± 0.1 Hz, mean ± s.e.m.; *P* = 8 × 10^−22^, two-sample t-test). For SST-Cre/R1117X^+/+^: *n* = 126 boutons from 4 animals. For SST-Cre, *n* = 184 boutons from 4 animals. * *P* < 0.05; ** *P* < 0.01; *** *P* < 0.001; n.s., not significant.

### Reduced NMDAR-mediated currents in SST interneurons in R1117X mice

The *in vivo* measurements confirmed SST interneurons as a probable culprit. It followed then that if SST interneurons were affected directly by the nonsense *Shank3* mutation, we should find reduced NMDAR signaling in the cell type. To demonstrate this link, we performed whole-cell voltage-clamp recordings from SST interneurons in Cg1/M2 in acute coronal brain slices (**Fig. 4a, b**). SST interneurons were identified based on fluorescence, because the SST-Cre and SST-Cre/R1117X^+/+^ animals were injected with AAV encoding Cre-dependent tdTomato and mTagBFP2. We placed a field stimulation electrode in layer 2/3 approximately 200 μm away from the patch electrode, in order to evoke postsynaptic currents in the patched SST interneuron. To isolate the NMDAR-mediated component of the excitatory postsynaptic current (NMDAR EPSC), NBQX and picrotoxin were added to the bath solution. We found that SST interneurons in SST-Cre/R1117X^+/+^ mice have lower NMDAR EPSC, relative to SST-Cre littermates (**Fig. 4c, e, j**; n = 17 – 21 cells from 4 – 5 mice for each condition). To determine the contribution of NMDAR subunits, we added the GluN2B-specific blocker ifenprodil. We specifically targeted GluN2B, because cortical SST interneurons contain an abundance of GluN2B transcripts [38, 53], and the subunit contributes a considerable amount of NMDAR-mediated current in the related low-threshold spiking interneuron subtype in the rodent prefrontal cortex [54]. In SST-Cre/R1117X^+/+^ animals, we found that GluN2B constituted a smaller fraction of the total NMDAR EPSC, compared to SST-Cre controls (**Fig. 4d, f, k**; n = 13 – 21 cells from 4 – 5 mice for each condition). Furthermore, there was no detectable effect of the R1117X mutation on the voltage activation curve, suggesting that changes in the mutants were not due to differences in the channel properties of NMDAR (**Fig. 4l**; n = 12 – 23 cells from 4 – 5 mice for each condition). We also measured pyramidal neurons (**Fig. 4m**), and they too had reduced NMDAR EPSC (**Fig. 4n, o**), although with no change in the GluN2B proportion (**Fig. 4n, p**), in animals with the R1117X mutation. Overall, these results demonstrate that the nonsense *Shank3* mutation strongly diminishes NMDAR currents in prefrontal SST interneurons.

**Fig. 4.**
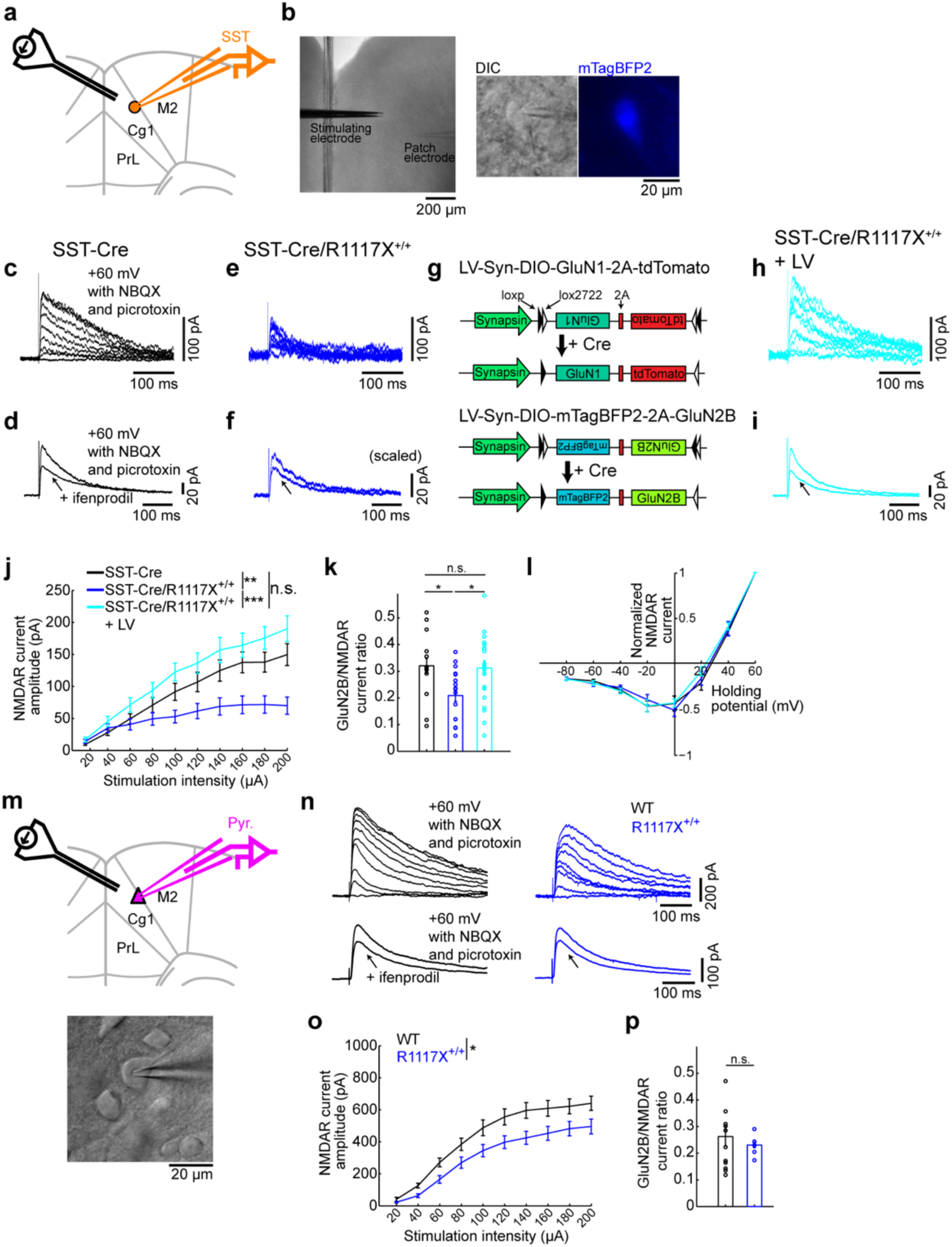
NMDAR currents in SST interneurons are diminished by the R1117X mutation in *Shank3*. (a) Schematic of the slice electrophysiology experiments for SST interneurons in Cg1/M2. (b) Images from an experiment. Left, wide-field differential interference contrast (DIC) image. Middle, a neuron and the patch pipette under DIC. Right, the same neuron identified as a SST interneuron based on mTagBFP2 fluorescence. (c) Example NMDAR-mediated excitatory postsynaptic currents (NMDAR EPSC) in response to stimulation intensities from 20 to 200 μA in steps of 20 μA for SST-Cre animals. (d) Example NMDAR EPSC without and with ifenprodil for SST-Cre animals. (e, f) Same as (c, d) but for SST-Cre/R1117X^+/+^ animals. (g) Schematic of LV-Syn-DIO-GluN1-2A-tdTomato for Cre-dependent GluN1 overexpression and LV-Syn-DIO-mTagBFP2-2A-GluN2B for Cre-dependent GluN2B overexpression. (h, i) Same as (c, d) but for SST-Cre/R1117X^+/+^ animals with lentiviral-mediated overexpression of GluN1 and GluN2B in SST interneurons (LV). (j) NMDA EPSC amplitude as a function of stimulation intensity. Two-way ANOVA with genotype (SST-Cre, SST-Cre/R1117X^+/+^ and LV) as between-subjects factor and stimulation intensity as within-subjects factor revealed a significant interaction between the two factors (F(18) = 10.1, *P* = 4 × 10^−24^). For SST-Cre, *n* = 18 cells, 5 mice. For SST-Cre/R1117X^+/+^, *n* = 21 cells, 4 mice. For LV, *n* = 17 cells, 5 mice. (k) GluN2B to NMDAR EPSC ratio. One way ANOVA with genotype as between-subjects factor revealed a significant main effect (F(2) = 4.5, *P* = 0.02). Post-hoc Tukey-Kramer tests, SST-Cre (0.32 ± 0.04, mean ± s.e.m.) versus SST-Cre/R1117X^+/+^ (0.21 ± 0.02) *P* = 0.04; SST-Cre versus LV (0.31 ± 0.03), *P* = 0.98; SST-Cre/R1117X^+/+^ versus LV, *P* = 0.03. For SST-Cre, *n* = 13 cells, 4 mice. For SST-Cre/R1117X^+/+^, *n* = 17 cells, 5 mice. For LV, *n* = 21 cells, 5 mice. (l) Normalized NMDAR EPSC as a function of holding potential. Two-way ANOVA with genotype as between-subjects factor and holding potential as within-subjects factor did not detect a main effect of genotype (F(2) = 0.12, *P* = 0.89), nor an interaction between the two factors (F(14) = 0.68 *P* = 0.80). For SST-Cre, *n* = 12 cells, 4 mice. For SST-Cre/R1117X^+/+^, *n* = 19 cells, 5 mice. For LV, *n* = 23 cells, 5 mice. (m) Schematic of the slice electrophysiology experiments for pyramidal neurons in Cg1/M2. A DIC image from an experiment. (n) Same as (c) and (d) but for pyramidal neurons in wild type and R1117X^+/+^ animals. (o) Same as (j) but for pyramidal neurons. Two-way ANOVA with genotype (wild type and R1117X^+/+^) as between-subjects factor and stimulation intensity as within-subjects factor revealed a significant interaction between the two factors (F(9) = 2.0, *P* = 0.04). For wild type, *n* = 28 cells, 4 mice. For R1117X^+/+^, *n* = 17 cells, 5 mice. (p) Same as (k) but for pyramidal neurons. Two-tailed t-test did not detect any difference (*P* = 0.52) between wild type (0.27 ± 0.04) and R1117X^+/+^ (0.23 ± 0.01). For wild type, *n* = 12 cells, 3 mice. For R1117X^+/+^, *n* = 7 cells, 2 mice. * *P* < 0.05; ** *P* < 0.01; *** *P* < 0.001; n.s., not significant.

### GluN2B upregulation in SST interneurons reverses *in vitro* and *in vivo* phenotypes in R1117X mice

We wanted to devise an approach to ameliorate the deficits in SST interneurons in animals with the R1117X mutation. There are a couple of clues from the electrophysiological data to suggest NMDAR upregulation as a viable approach. One clue is that the NMDAR EPSC was reduced but not eliminated by the R1117X mutation, indicating that there are functional receptor complexes. For example, other members of the SHANK protein family also associate with glutamatergic receptors, and can be responsive to NMDAR upregulation. Another clue is the decreased contribution of GluN2B to NMDA EPSC in SST interneurons in SST-Cre/R1117X^+/+^ animals. The preferential reduction suggests that selective restoration of GluN2B-containing NMDARs may be effective. To this end, we designed lentiviral vectors to overexpress NMDAR subunits in a Cre-dependent manner (**Fig. 4g, Supplementary Fig. 4**). Because it was unclear which subunit is the factor limiting the number of functional GluN2B-containing NMDARs at synapses, we introduced both GluN1 and GluN2B subunits into the medial prefrontal cortex (Cg1/M2) of SST-Cre/R1117X^+/+^ animals. The cell-type-specific lentiviral manipulation enhanced the evoked NMDAR currents (**Fig. 4h, j**) and increased the relative contribution of GluN2B (**Fig. 4i, k**) in SST interneurons, without altering the NMDAR activation curve (**Fig. 4l**). The results suggest that upregulation of GluN2B-containing NMDARs is effective at reversing the deficient NMDAR currents in SST interneurons associated with the R1117X mutation.

Encouraged by the success of the lentiviral manipulation *in vitro*, we expect that restoration of NMDAR currents in SST interneurons may correct the *in vivo* phenotype of aberrant synaptic calcium dynamics in R1117X^+/+^ animals. In these experiments, we first applied lentiviral manipulation and AAV for GCaMP expression, and then performed *in vivo* imaging three weeks later to characterize calcium transients in SST interneurons and pyramidal dendritic spines (**Fig. 5a**). Under the two-photon microscope, we found cells expressing three fluorophores *in vivo*, indicating GCaMP6s-expressing SST interneurons with GluN1 and GluN2B overexpression (**Fig. 5b**). Post-hoc histology showed that at the injection site, 63% of the SST interneurons received both lentiviruses (**Fig. 5c**). In terms of neural dynamics *in vivo*, the SST interneuron-specific lentiviral manipulation elevated SST interneuron activity (2.5 ± 0.2 Hz, mean ± s.e.m.; compare with SST-Cre/R1117X^+/+^: 0.97 ± 0.08 Hz; *P* = 2 × 10^−6^, post-hoc Tukey-Kramer test after one-way ANOVA test; **Fig. 5d**), and restored the calcium event rate in dendritic spines to levels comparable to control animals (0.79 ± 0.03 Hz, mean ± s.e.m.; compare with wild type: 0.83 ± 0.01 Hz; *P* = 0.52, post-hoc Tukey-Kramer test after one-way ANOVA test; **Fig. 5e**). Altogether, the *in vivo* causal manipulations confirmed deficient dendritic inhibition by SST interneurons as a microcircuit mechanism underlying the excessive spine calcium signals.

**Fig. 5.**
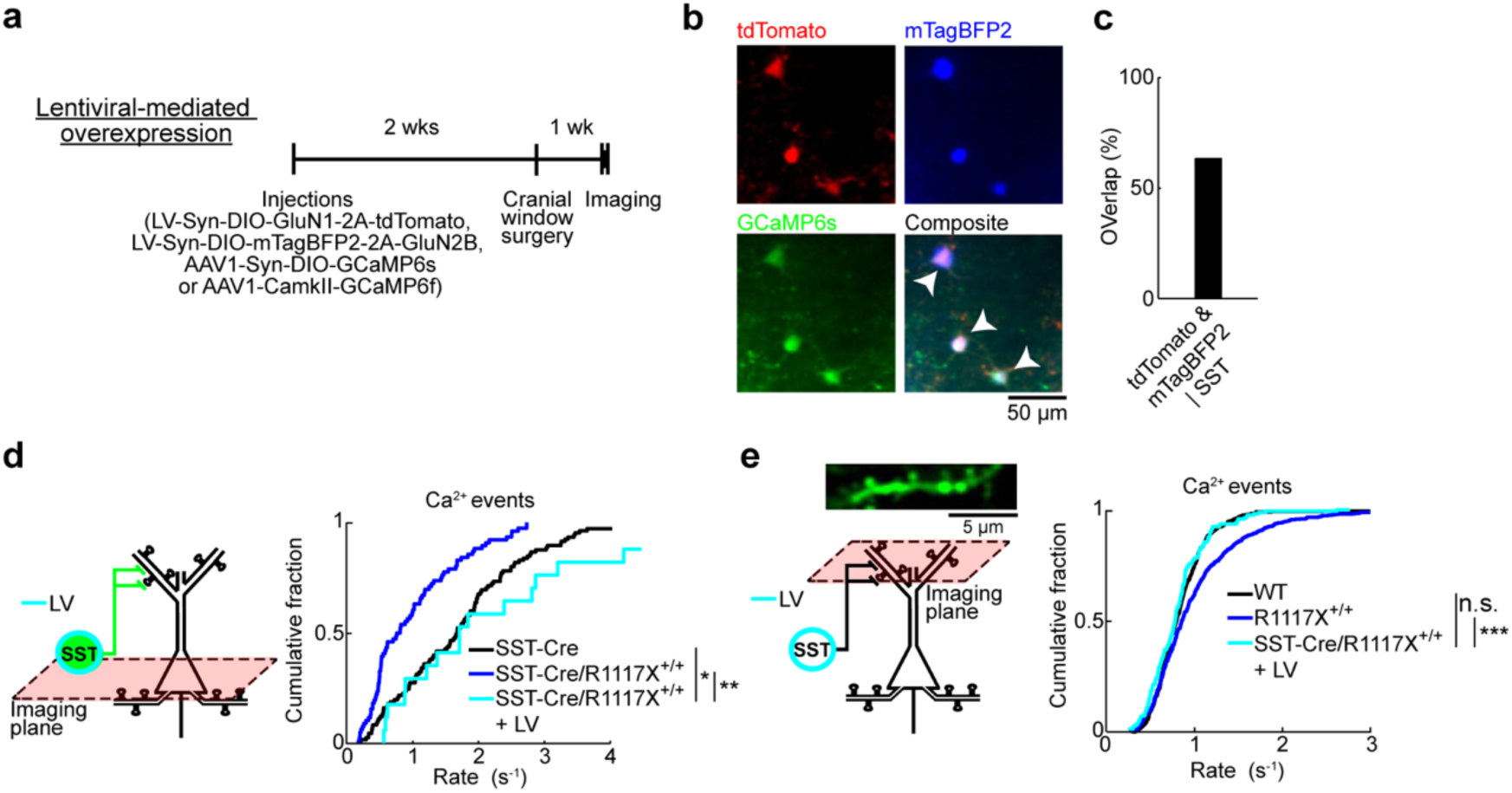
Elevating SST interneuron activity *in vivo* ameliorates aberrant spine calcium dynamics. (a) Top, timeline for experiments involving lentiviral-mediated overexpression of GluN1 and GluN2B in SST interneurons in SST-Cre/R1117X^+/+^ animals (LV). (b) *In vivo* two-photon images of tdTomato (red), mTagBFP2 (blue), and GCaMP6s (green). Arrowhead, cell that expressed the three transgenes. (c) Histological quantification of efficiency of lentiviral-mediated transduction of SST interneurons (tdTomato & mTagBFP2 | SST, 63%). *n* = 219 cells, 3 mice. (d) For SST interneurons, cumulative plot of the rate of calcium events (LV: 2.5 ± 0.2 Hz, mean ± s.e.m.). SST-Cre (black line) and SST-Cre/R1117X^+/+^ (blue line) were duplicated from Fig. 3e for comparison (one-way ANOVA with genotype (SST-Cre, SST-Cre/R1117X^+/+^ and LV) as the between-subjects factor: F(2,196) = 16.2, *P* = 3 × 10^−7^; post-hoc Tukey-Kramer tests for SST-Cre versus LV: *P* = 0.02, SST-Cre/R1117X^+/+^ versus LV: *P* = 2 × 10^−6^). For LV, *n* = 18 cells from 3 animals. (e) For apical dendritic spines in layer 1, cumulative plot of the rate of calcium events (LV: 0.79 ± 0.03 Hz, mean ± s.e.m.). Wild type (black line) and R1117X^+/+^ (blue line) duplicated from Fig. 2c for comparison (one-way ANOVA with experiment type as the between-subjects factor: F(2,1279) = 23.5, *P* = 1 × 10^−10^; post-hoc Tukey-Kramer tests for wild type versus LV: *P* = 0.52, R1117X^+/+^ versus LV: *P* = 6 × 10^−7^). For LV, *n* = 171 spines from 3 animals. * *P* < 0.05; ** *P* < 0.01; *** *P* < 0.001; n.s., not significant.

**Fig. 6.**
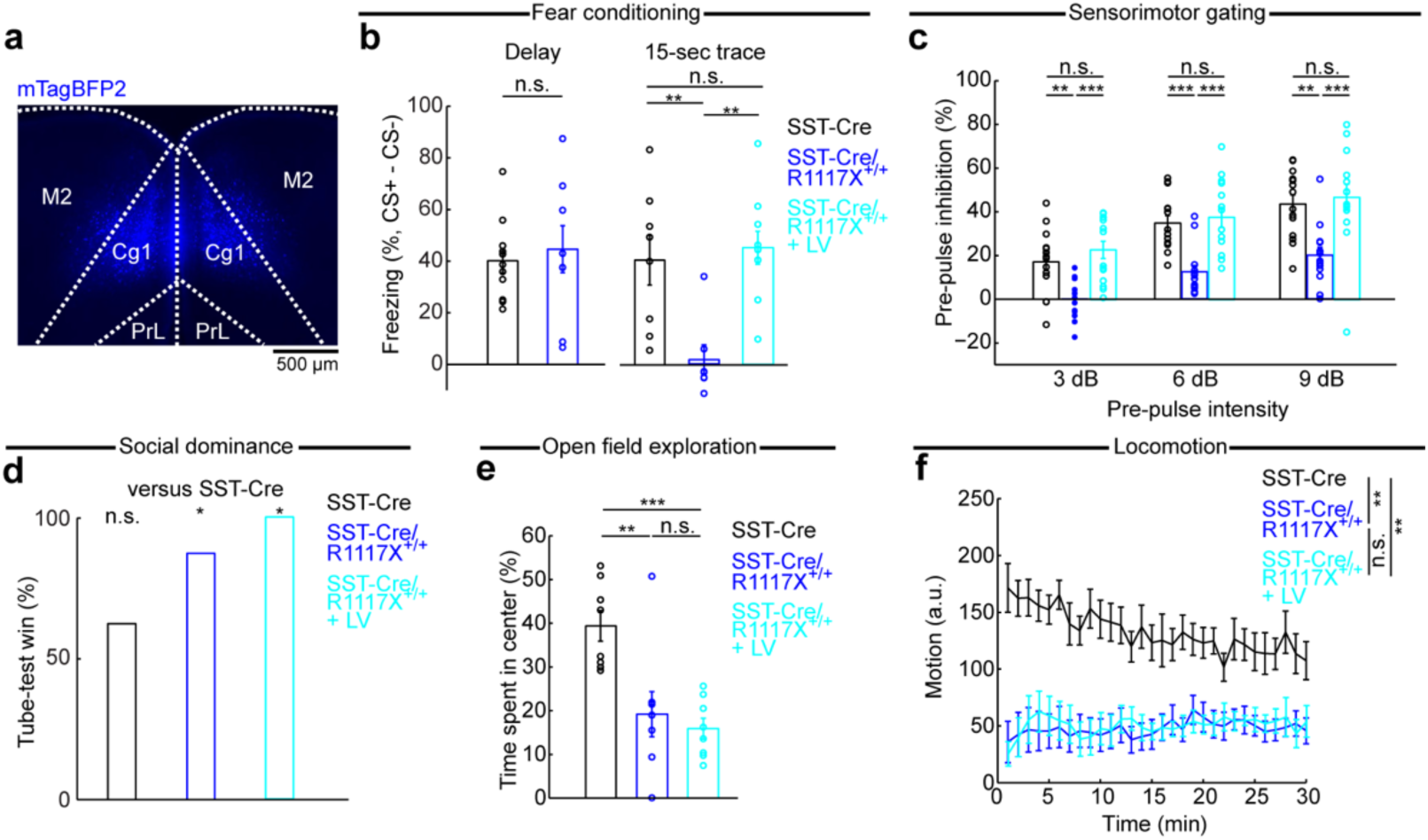
Behavioral consequences of manipulating prefrontal cortical SST interneurons. (a) Coronal histological section, showing the extent of mTagBFP2 expression in SST-Cre/R1117X^+/+^ animals with bilateral, lentiviral-mediated overexpression of GluN1 and GluN2B in SST interneurons (LV). (b) Fraction of time spent freezing during CS+ and trace period (if any), subtracted by that during CS- and trace period (if any), on the testing day. Freezing behavior in delay fear conditioning (SST-Cre: 47 ± 6%, SST-Cre/R1117X^+/+^: 50 ± 5%, mean ± s.e.m.; *P* = 0.7, Wilcoxon rank-sum test; *n* = 8 and 12 animals for SST-Cre and R1117X^+/+^ respectively), and trace fear conditioning (SST-Cre: 41 ± 10%, SST-Cre/R1117X^+/+^: 2 ± 6%, LV: 45 ± 6%, mean ± s.e.m.). LV restored trace fear learning in SST-Cre/R1117X^+/+^ (SST-Cre versus SST-Cre/R1117X^+/+^, *P* = 0.004; SST-Cre versus LV, *P* = 0.76; SST-Cre/R1117X^+/+^ versus LV, *P* = 4 × 10^−4^; Wilcoxon rank-sum test). *n* = 8 animals for SST-Cre, 7 for SST-Cre/R1117X^+/+^ and 10 for LV. (c) Pre-pulse inhibition as a measure of sensorimotor gating. Two-way mixed ANOVA with genotype (SST-Cre, SST-Cre/R1117X^+/+^, LV) as between-subjects factor and pre-pulse intensity (3, 6, 9 dB) as within-subjects factor found significant main effects of genotype (F(2) = 17.9, *P* = 3 × 10^−6^) and pre-pulse intensity (F(2) = 43.5, *P* = 6 × 10^−12^), but a non-significant interaction (F(4) = 0.4, *P* = 0.78). One-way ANOVA with genotype as a between-subjects factor was followed by post-hoc Tukey-Kramer’s tests to determined statistical significance for each of the pre-pulse intensity levels. *n* = 14 animals for each genotype. (d) Percentage of wins for each genotype, against SST-Cre animals: SST-Cre (62.5%; *P* = 0.36, binomial test), SST-Cre/R1117X^+/+^ (87.5%; *P* = 0.04), LV (100%). *n* = 8 animals for each genotype. (e) Percentage of time spent in center of an open field: SST-Cre (40 ± 3%) versus SST-Cre/R1117X^+/+^ (19 ± 5%), *P* = 0.005; SST-Cre versus LV (16 ± 2%), *P* = 2 × 10^−4^; SST-Cre/R1117X^+/+^ versus LV, *P* = 0.88. Wilcoxon rank-sum tests. *n* = 8 animals for each genotype. (f) Motion (mean ± s.e.m.) as a function of time in an open field. Motion averaged over the 30-min period: SST-Cre (133 ± 13, arbitrary units (a.u.)) versus SST-Cre/R1117X^+/+^ (48 ± 9), *P* = 0.001; SST-Cre versus LV (50 ± 10), *P* = 0.001; SST-Cre/R1117X^+/+^ versus LV, *P* = 0.88. All Wilcoxon rank-sum tests. *n* = 8 animals for each genotype. * *P* < 0.05; ** *P* < 0.01; *** *P* < 0.001; n.s., not significant.

### Enhancing dendritic inhibition reestablishes prefrontal-dependent learning in R1117X mice

Next, we asked to what extent dysfunction of the dendritic inhibition mechanism may influence behavior. In trace fear conditioning, an auditory stimulus (conditioned stimulus, CS+) is paired with a footshock (unconditioned stimulus, US) with an intervening trace period. This assay requires associative learning across a temporal gap, and therefore is a classic task requiring an intact medial prefrontal cortex [55]. SST-Cre/R1117X^+/+^ animals did not freeze after conditioning with a 15-s trace period, but were sensitive to delay fear conditioning in which the CS co-terminated with the US, suggesting an impairment specific to temporal association and not learning in general (trace: 2 ± 6%, delay: 50 ± 5%, mean ± s.e.m.; **Fig. 6b**). Lentiviral-mediated overexpression of GluN1 and GluN2B in SST interneurons, applied to the medial prefrontal cortex (Cg1/M2) bilaterally (**Fig. 6a, b**), returned trace fear learning to levels comparable to control littermates (45 ± 6%; compare to SST-Cre controls: 47 ± 6%; SST-Cre/R1117X^+/+^ versus SST-Cre: *P* = 0.004, SST-Cre/R1117X^+/+^ with lentivirus versus SST-Cre: *P* = 0.76, Wilcoxon rank-sum test; **Fig. 6b**). These results indicate that the deleterious impact of the nonsense *Shank3* mutation on prefrontal-dependent learning relies on perturbing dendritic inhibition.

### Effects of restoring dendritic inhibition on social-, anxiety-, and locomotor-related behaviors in R1117X mice

We additionally characterized behaviors using 4 assays: prepulse inhibition (PPI), social dominance, open field exploration, and locomotion. These assays were selected because a previous study found alterations in the same or similar behavioral tests in mice with the R1117X mutation in *Shank3* [26]. Here, the goal was to test whether upregulating NMDAR expression in prefrontal SST interneurons can alleviate the behavioral deficits. PPI, a measure of sensorimotor gating, is impaired in schizophrenia-spectrum patients and related animal models [56]. Adolescent SST-Cre/R1117X^+/+^ mice had deficient PPI compared to SST-Cre controls (**Fig. 6c**). Bilateral lentiviral-mediated overexpression of GluN1 and GluN2B in SST interneurons in Cg1/M2 was sufficient to restore PPI for the SST-Cre/R1117X^+/+^ mice (**Fig. 6c**). However, not all behaviors were affected by the prefrontal lentiviral manipulation. For social dominance, we adapted the tube test in which two animals faced each other inside a narrow tube and losing was defined by retreat [40]. Against SST-Cre controls, the SST-Cre/R1117X^+/+^ animals won frequently, and the dominance remained after lentiviral manipulation (**Fig. 6d**). Moreover, SST-Cre/R1117X^+/+^ mice were less likely to spend time in the center of an open-field chamber, an indicator of anxiety-like behavior (**Fig. 6e**). This behavioral phenotype persisted after lentiviral manipulation (**Fig. 6e**). Hypo-locomotion in SST-Cre/R1117X^+/+^ mice was similarly resistant to lentiviral manipulation (**Fig. 6f**). Overall, of behaviors examined, trace fear learning and sensorimotor gating responded to the lentiviral manipulation, suggesting that the deleterious effects of the nonsense *Shank3* mutation on those behaviors depended on SST interneuron-mediated dendritic inhibition in prefrontal cortex. By contrast, other social-, anxiety-, and locomotor-related behavior deficits induced by the R1117X mutation appear to be mediated by other mechanisms or brain regions.

## Discussion

By imaging calcium transients in apical dendritic spines in the mouse medial prefrontal cortex (Cg1/M2) *in vivo*, we have provided a direct glimpse of the dysfunctional synapses in an awake brain harboring a nonsense *Shank3* mutation. There are two major results. First, we showed an abnormal elevation of synaptically evoked calcium influx in dendritic spines of pyramidal neurons. Second, we identified the loss of inhibitory control on dendritic spines as a microcircuit mechanism responsible for the synaptic alterations with behavioral consequences. Specifically, we showed that targeting GluN2B receptors in dendrite-targeting interneurons was a promising approach to correct the synaptic and behavioral deficits for the mutant mice.

The abnormal increases in calcium transients in the dendritic spines were unexpected. During normal synaptic activation, the dendritic spine is depolarized and calcium ions enter primarily through the opening of voltage-gated NMDARs [57, 58]. Disruption of SHANK3 has been reported to weaken the synaptic activation, typically measured by counting functional NMDARs [12, 59] and recording postsynaptic potentials [13]. R1117X could be a neomorphic mutation, although previous studies have provided biochemical data to suggest that it is a partial loss-of-function mutation [9, 26]. Therefore, the direct impact of reduced SHANK3 function on dendritic spines should reduce synaptic calcium influx. Instead, we observed increased synaptic calcium signals, and our results indicate a two-step mechanism: effect of mutation on dendrite-targeting SST GABAergic neurons which in turn influences calcium signals at the dendritic spines. Structurally, SST interneurons synapse onto both spine heads and dendritic shafts, and are thus well positioned to control the excitability of single or cluster of dendritic spines [49, 50, 60]. Functionally, experiments involving optogenetic stimulation of SST interneurons in brain slices have demonstrated that these inhibitory inputs powerfully modulate calcium influx in dendritic spines [50, 51]. SST interneurons likely have a more prominent role in awake animals relative to *in vitro* conditions, because SST interneurons are modulated by behavioral state and are particularly active during quiet wakefulness *in vivo* [61]. Alterations that reduce the activity of SST interneurons could therefore put dendritic spines of pyramidal neurons at a more excitable state, raising the amount of calcium influx during synaptic activation. Results from multiple approaches in our study, including imaging the activity of SST interneurons and monitoring synaptic calcium transients while manipulating dendritic inhibition, are in full agreement with this local-circuit scheme.

A number of studies of the neocortex or hippocampus have revealed diminished GABAergic signaling in mouse models with defective SHANK proteins [20-22]. However, most of the evidence involved characterizing inhibitory currents in the soma or counting inhibitory synaptic markers, which do not allow for the identification of the contributing GABAergic cell type. Our study adds to this body of growing evidence that GABAergic neurons are implicated in SHANK3 pathology, and specifically highlights the dendrite-targeting SST interneurons. In terms of understanding how excitation-inhibition imbalance may underlie neurodevelopmental disorders [62-64], the potential involvement of heterogeneous GABAergic cell types is an important point as it emphasizes the complex ways by which inhibition can shape cortical activity [65], and how imbalance could occur at specific neuronal compartments such as the dendrites. Therefore, our results highlight the need for more studies that can examine the impact of cell-type-specific perturbation of *Shank3* [66]. The loss of dendritic inhibition may interact with other forms of SHANK3-related channelopathy affecting dendritic excitability [67] or metabotropic glutamate receptor signaling [11, 68], which should be tested in future studies.

In experiments involving causal manipulations, we focused on GluN2B-containing NMDARs. We picked this molecular target for biological reasons, because not only is SHANK3 a well-established scaffold for NMDARs at postsynaptic sites [1], but specifically SST interneurons abundantly express GluN2B transcripts [38, 53] and carry significant GluN2B-mediated currents [54]. We should note that there are also substantial translational implications for testing this particular NMDAR subunit. For ASD, the *GRIN2B* gene that encodes GluN2B is among a handful of top-scoring, high-confidence risk gene in the database curated by the Simons Foundation Autism Research Initiative, based on a large number of genetic studies [69-71]. Intriguingly, akin to how *SHANK3* is linked to multiple neurodevelopmental disorders, *GRIN2B* mutations have also been associated with intellectual disability and schizophrenia [72, 73]. The genetic underpinnings, as well as some promising results in pharmacological tests, have led to the suggestion that GluN2B-specific positive allosteric modulators may have therapeutic potential [74]. In animals, other studies have manipulated the firing of SST interneurons to modulate pyramidal neuron activity [75, 76]. Our study differs by focusing on the outcomes at the synaptic level, and hints at the possibility that augmenting NMDAR signaling, if done with cell-type and subunit specificity, may be a promising strategy for correcting the neural circuit dysfunction.

What are the implications for the elevated synaptic calcium signals? Although there are many reports of synaptic pathology associated with neurodevelopmental disorders, few studies have directly looked at functional calcium signals in dendritic spines *in vivo*. For dendritic spines, local calcium homeostasis is a tightly regulated process. Calcium entry is restricted spatially, as the spine head acts as an electrically isolated compartment [77]. Calcium is also controlled temporally, as the ions interact with other postsynaptic molecules over a time scale from seconds to minutes [78]. The spread of the calcium signal orchestrates the molecular processes for activity-dependent plasticity, and has been shown to be necessary and sufficient for synaptic potentiation [79, 80]. Therefore, dysregulation of the calcium signaling in dendritic spines, particularly during the adolescence period, is expected to hinder the normal maturation of the prefrontal cortical circuitry. Moreover, by showing that calcium is excessive rather than missing, our results hint at the possibility that the problem for the mutants may not be a lack of plasticity, but rather maladaptive plasticity [63].

The maintenance of information in the prefrontal cortex requires reverberation of neural activity. Computational models of cortical microcircuits that include pyramidal neurons and multiple interneurons subtypes have illustrated elegantly the role of the dendrite-targeting cell type in filtering out distractor stimuli and thereby stabilizing the information during the delay period associated with short-term memory [81]. This role in maintenance is supported by electrophysiological recordings and optogenetic manipulation of prefrontal SST interneurons, which exhibit delay-period specificity in its neural activity and behavioral influences [82, 83]. Prior studies of normal behavior, thus, predict a deficit in short-term memory behaviors associated with abnormal distractor-filtering circuitry adversely impacted by SHANK3 dysfunction. We tested this prediction using trace fear conditioning in which the animal has to associate the footshock to a preceding sensory stimulus that is no longer present, thus requiring the short-term maintenance of sensory information. Fully consistent with the prediction, we found impairment in trace fear learning in the R1117X mutants, which could be rescued by restoring deficient dendritic inhibition. The same dendritic inhibition mechanism has also been posited to be involved in gating irrelevant inputs during context-dependent decisions [26], therefore one prediction is that these R1117X mutants may also be impaired in more cognitive demanding tasks involving behavioral flexibility [84].

The circuit-level consequence of a *Shank3* mutation will vary depending on the nature of the mutation. Notably, we showed that *Shank3B*^+/-^ mice had diminished synaptic calcium transients in the apical dendritic spines, which was opposite to what we observed for R1117X^+/+^ animals. These global *Shank3* mutations affect all cell types in the cortical microcircuit. Although the direct effect of diminished NMDAR function on pyramidal neurons is reduced synaptic excitability, there is also an indirect effect of disinhibition via diminished SST interneuron firing. Indeed, the slice electrophysiology results demonstrated that the missense *Shank3* mutation diminishes NMDAR-mediated transmission in both pyramidal cells (**Fig. 4o**) and SST interneurons (**Fig. 4j**). These direct and indirect effects are expected to compete – suppressing and promoting dendritic calcium signals, respectively. It is known that knockout of *Shank3B* substantially reduces the expression of SHANK3, including complete elimination of the SHANK3_α_ and SHANK3_β_ isoforms [10]. Our interpretation of the data is that NMDAR function is impaired in *Shank3B*^+/-^ mice, with a predominant direct effect that suppresses synaptic calcium transients. By contrast, the partial loss-of-function induced by the R1117X point mutation appears to be milder, as truncated SHANK3 remains in postsynaptic densities [26]. In this case, potentially with other unknown reason that biases the mutation towards affecting NMDAR in SST interneurons, the balance of the circuit interaction tilts towards dendritic disinhibition and consequently synaptic calcium transients are elevated.

In conclusion, we initiated this study because the R1117X mutation in *SHANK3* has been associated with unusual behavioral disturbances, yet its impacts on neural circuits were not understood. Here, we show that the nonsense *Shank3* mutation leads to aberrant synaptic calcium transients in pyramidal neurons *in vivo*, mediated through deficient dendritic inhibition in the mouse prefrontal cortex. It is an example for how a single genetic alteration has the potential to affect different cell types in a neural circuit, and it is the balance of the circuit-level interactions that would dictate the overall impact on neural function and behavior.

## Acknowledgements

We thank K. Kwan and M. Picciotto for comments on the initial version of this paper; C.S. Chan for advice on electrophysiology; M. Picciotto for supplying pAAV-CMV-dsRed-pSico-GFP; S. Tomita for discussions; I.-J. Kim for use of microscope. This work was supported by Simons Foundation Autism Research Initiative Pilot Award (A.C.K.). Additional support was provided by National Institute of Mental Health grants R01MH112750 (A.C.K.) and R21MH110712 (A.C.K.), NARSAD Young Investigator Grant (A.C.K.), Alzheimer’s Association Research Fellowship AARF-17-504924 (F.A.), and James Hudson Brown-Alexander Brown Coxe Postdoctoral Fellowship (F.A.).

## Author contributions

F.A. and A.C.K. designed the research. F.A. performed surgery, imaging, behavioral experiments, and histology, and analyzed the data. L.-X.S. and F.A. performed the slice electrophysiology experiments and analyzed the data. D.M.G., S.P., R.S.D. performed molecular cloning, viral packaging, and *in vitro* validation. K.S. assisted with the imaging data analysis. C.P. assisted with PPI experiments. F.A. and A.C.K wrote the paper, with input from all other authors.

## Data availability

The data that support the findings of this study are available from the corresponding author upon reasonable request.

## Code availability

MATLAB scripts for the present study are available from the corresponding author upon reasonable request.

## Supplementary Figures

**Supplementary Fig. 1.**
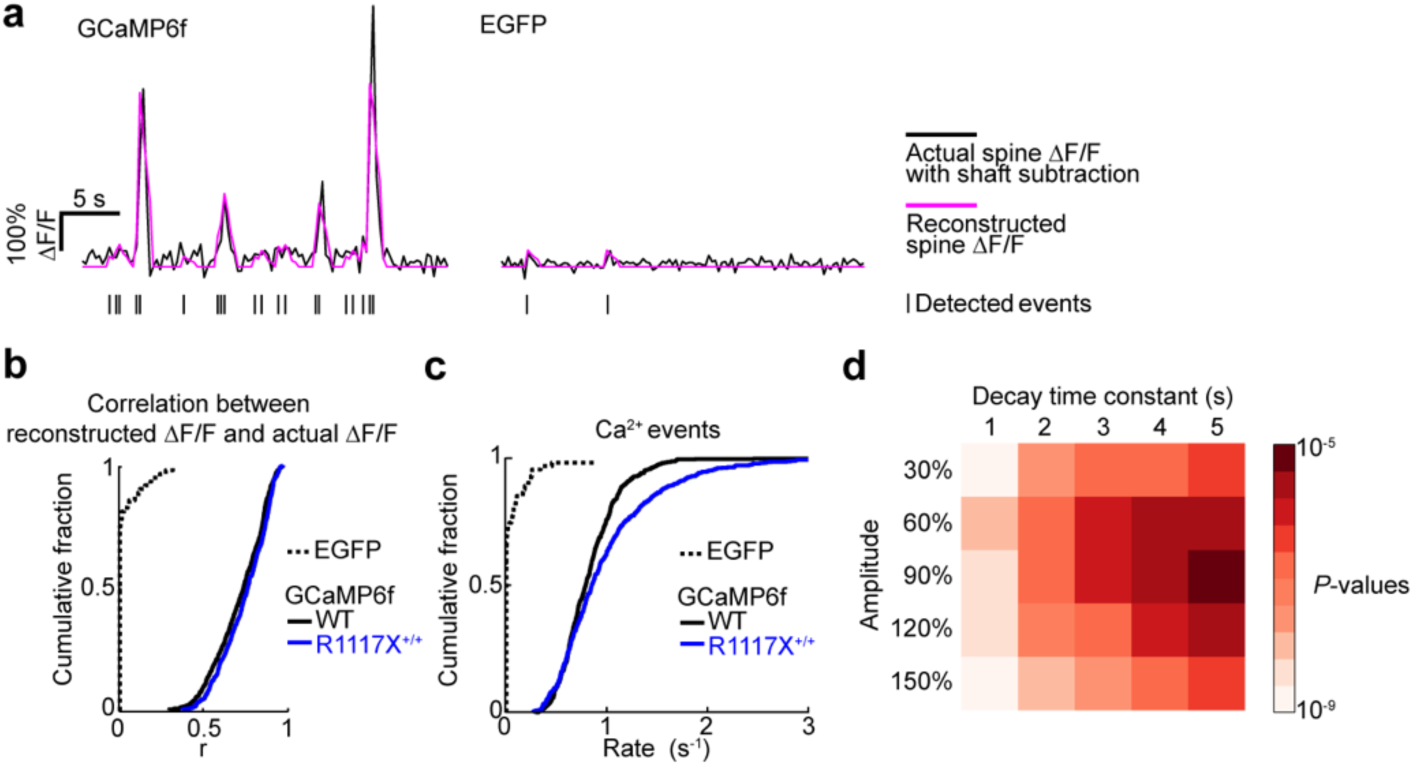
Additional analyses of synaptic calcium transients in dendritic spines. (a) Left, example segment of shaft-subtracted spine signal (black) and reconstructed spine signal (magenta) for GCaMP6f. The reconstructed spine signal was based on a peeling algorithm to detect calcium events (tick marks) – synaptic calcium signals that match an elementary template. The peeling algorithm follows Lutcke H et al., Front. Neural Circuits, 7, 201 (2013). For details, see Materials and Methods. Right, same as left but for EGFP, a non-calcium-dependent fluorophore. (b) Summary plot of the correlation between actual and reconstructed ΔF/F at imaging frames in which a calcium event was detected (EGFP: 0.02 ± 0.01, GCaMP6f in R1117X^+/+^: 0.739 ± 0.006, GCaMP6f in wild type: 0.712 ± 0.006, mean ± s.e.m). For EGFP, n = 76 spines from 3 animals. For GCaMP6f in R1117X^+/+^, n = 553 dendritic spines from 4 animals. For GCaMP6f in wild type, n = 558 dendritic spines from 5 animals. (c) Cumulative plot of the rate of detected events (EGFP: 0.05 ± 0.01 Hz). Wild type (black line) and R1117X^+/+^ (blue line) duplicated from Fig. 2c for comparison. For EGFP, n = 76 spines from 3 animals. (d) Left, *p*-values of two-sample t-tests comparing spine calcium events in R1117X^+/+^ to WT as a function of template parameters. * *P* < 0.05; ** *P* < 0.01; *** *P* < 0.001; n.s., not significant.

**Supplementary Fig 2.**
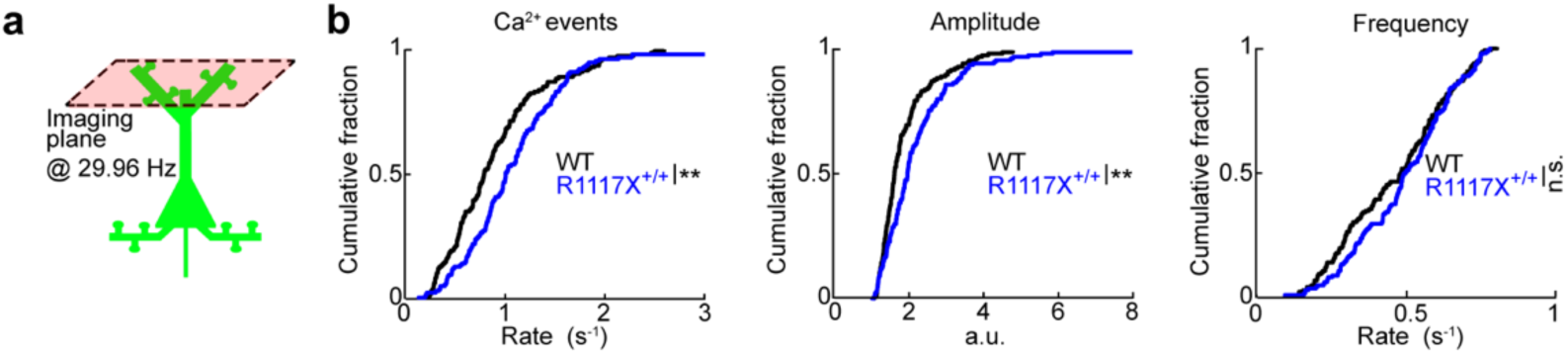
Imaging dendritic spines at higher frame rates. (a) Schematic of apical dendritic spines in layer 1 at 29.96 Hz using a resonant-based scanner. (b) Left, cumulative plot of the rate of calcium events using template-matching and peeling (R1117X^+/+^: 1.08 ± 0.04 Hz, wild type: 0.84 ± 0.04 Hz, mean ± s.e.m.; *P* = 0.002, two-sample t-test). After a binning procedure (see Materials and methods): middle, cumulative plots of amplitude (R1117X^+/+^: 2.22 ± 0.09, wild type: 1.87 ± 0.06; *P* = 0.001, two-sample t-test), and right, frequency of binned calcium events (R1117X^+/+^: 0.50 ± 0.01, wild type: 0.47 ± 0.01; *P* = 0.12). For R1117X^+/+^, *n* = 154 dendritic spines from 4 animals. For wild type, *n* = 154 dendritic spines from 4 animals. * *P* < 0.05; ** *P* < 0.01; *** *P* < 0.001; n.s., not significant.

**Supplementary Fig 3.**
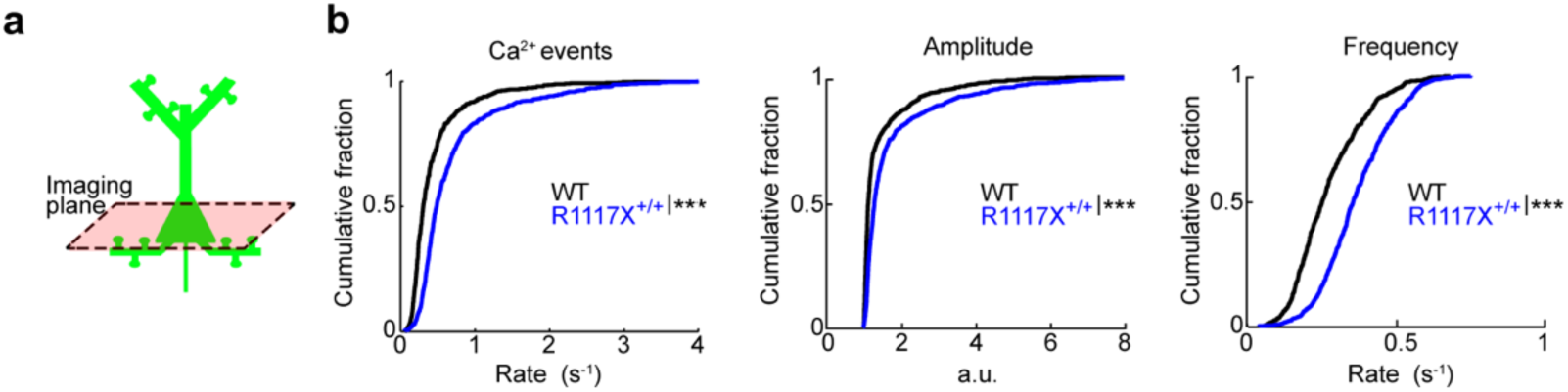
Impact of R1117X^+/+^ on pyramidal neuron cell body activity. (a) Schematic of layer 2/3 pyramidal neuron cell body imaging. (b) Left, cumulative plot of the rate of calcium events using template-matching and peeling (R1117X^+/+^: 0.72 ± 0.02 Hz, wild type: 0.44 ± 0.02 Hz, mean ± s.e.m.; *P* = 1 × 10^−19^, two-sample t-test). After a binning procedure (see Materials and methods): middle, cumulative plots of amplitude (R1117X^+/+^: 1.79 ± 0.04, wild type: 1.47 ± 0.03; *P* = 1 × 10^−10^, two-sample t-test), and right, frequency of binned calcium events (R1117X^+/+^: 0.372 ± 0.004 Hz, wild type: 0.279 ± 0.005 Hz.; *P* = 0.02, two-sample t-test). For R1117X^+/+^, *n* = 822 neurons from 5 animals. For wild type, *n* = 694 neurons from 4 animals. * *P* < 0.05; ** *P* < 0.01; *** *P* < 0.001; n.s., not significant.

**Supplementary Fig. 4.**
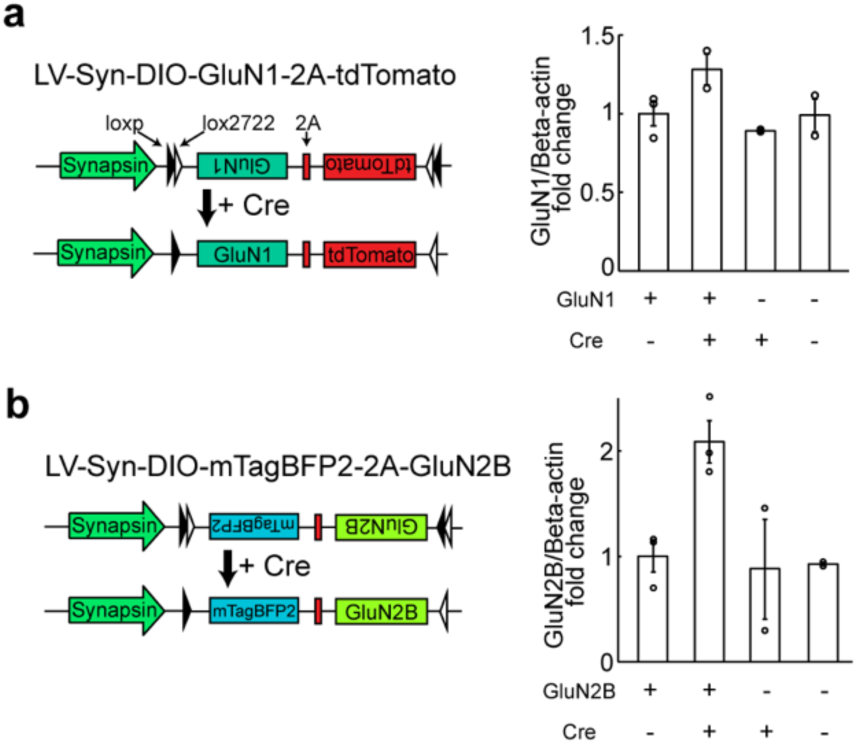
Validation of Cre-dependent overexpression of NMDA receptors. (a) Left, schematic of LV-Syn-DIO-GluN1-2A-tdTomato for Cre-dependent GluN1 overexpression. Right, protein expression levels via Western blot to validate Cre-dependent GluN1 overexpression. GluN1 signals relative to beta-actin signals for 4 conditions (all values normalized to the GluN1, no Cre condition): GluN1, no Cre (1.00 ± 0.08); GluN1, Cre (1.28 ± 0.12), no GluN1, Cre (0.89 ± 0.01); no GluN1, no Cre (0.99 ± 0.12). (b) Left, schematic of LV-Syn-DIO-mTagBFP2-2A-GluN2B for Cre-dependent GluN2B overexpression. Right, protein expression levels via Western blot to validate Cre-dependent GluN2B overexpression. GluN2B signals relative to beta-actin signals for 4 conditions (all values normalize to the GluN2B, no Cre condition): GluN2B, no Cre (1.00 ± 0.15); GluN2B, Cre (2.09 ± 0.2); no GluN2B, Cre (0.88 ± 0.58); no GluN2B, no Cre (0.93 ± 0.02).

